# Interpretable Deep Learning Model Reveals Subsequences of Various Functions for Long Non-coding RNA Identification

**DOI:** 10.1101/2022.02.11.479495

**Authors:** Rattaphon Lin, Duangdao Wichadakul

## Abstract

Long non-coding RNAs (lncRNAs) play crucial roles in many biological processes and are implicated in several diseases. With the next-generation sequencing technologies, substantial un-annotated transcripts have been discovered. Classifying unannotated transcripts using biological experiments is more time-consuming and expensive than computational approaches. Several tools for identifying long non-coding RNAs are available. These tools, however, did not explain which features in their tools contributed to the prediction results. Here, we present Xlnc1DCNN, a tool for distinguishing long non-coding RNAs (lncRNAs) from protein-coding transcripts (PCTs) using a one-dimensional convolutional neural network with prediction explanations. The evaluation results of the human test set showed that Xlnc1DCNN outperformed other state-of-the-art tools in terms of accuracy and F1-score. The explanation results revealed that lncRNA transcripts were mainly identified as sequences with no conserved regions or with a region of transmembrane helix while protein-coding transcripts were mostly classified by conserved protein domains or families. The explanation results also conveyed the probably inconsistent annotations among the public databases, lncRNA transcripts which contain protein domains or families, as well as protein-coding transcripts which are nonsense-mediated decay or processed transcripts. Xlnc1DCNN is freely available at https://github.com/cucpbioinfo/Xlnc1DCNN.

## 1. Introduction

Long non-coding RNAs (lncRNAs) are RNAs that are not translated into proteins and longer than 200 nucleotides. lncRNAs play important roles in many critical biological processes, including gene expression, gene regulation, gene silencing, chromatin remodelling, acting as molecular scaffolds, etc. [1–3], and have been implicated in human diseases such as cancers and diabetes [4–7]. The enhancements of next-generation sequencing technology, i.e., RNA Sequencing (RNA-Seq) [8,9] have led to numerous discoveries of unannotated transcripts. However, classifying the innumerable numbers of unclassified sequences using experimental approaches is time-consuming and expensive. In contrast, computational approaches are faster and more convenient.

Most of the existing computational approaches for classifying lncRNA and protein-coding transcripts used feature extraction methods to obtain training features, e.g., the upgraded version of Coding Potential Calculator (CPC2) [10], CNIT [11], PLEK [12], CPAT [13], FEELnc [14], RNAsamba [15], LncADeep [16], and lncRNA_Mdeep [17]. Most of them used similar features such as the Fickett and hexamer scores, ORF length, and then topped up with additional sequence and structural features. Moreover, none of them explained how the features contributed to the model prediction results.

The deep learning algorithms have become very popular, especially for a dataset with a large number of data points and data dimensions as the features will be learned by the algorithms themselves during the training. Many convolutional neural networks (CNNs), the 2D-CNNs, have been widely used for image classification and segmentation applications [18] because of their great capability for extracting features from input data. Recently, many applications such as speech recognition, ECG monitoring [19] started to use 1D-CNN instead of the traditional machine learning approaches. The applications on detecting irregular heartbeats [20–22] have shown that using only a simple 1D-CNN could achieve a high prediction accuracy without explicitly addressing and extracting features as inputs for the models.

While most complex black-box models (e.g., boosting tree algorithms, ensemble models, deep neural networks) typically provide better learning performance, they usually are uninterpretable. To understand how a complex model learns to differentiate things, explainable artificial intelligence (XAI) has recently become one of the popular topics aiming to interpret and explain machine learning or deep learning models [23]. Explainable AI is essential for users to understand and trust the model prediction results. It can help illustrate what the models perceive and explain how these perceptions can be mapped with the underlying knowledge of the human. Some of the favoured approaches to obtain an explanation from a complex black-box model are LIME [24] and SHAP [25]. LIME builds a local surrogate model to explain individual prediction. SHAP (<u>Sh</u>apley <u>A</u>dditive ex<u>P</u>lanations) introduced SHAP values representing the unified measure of feature importance together with SHAP value estimation methods. DeepSHAP [26] was built based on the connection between the original SHAP and DeepLIFT [27] to explain the deep learning model and further refined and extended with relative background distributions and stacks of mixed model types.

With still some ambiguities in classifying lncRNA and mRNA sequences based on training features, together with the promising results of 1D-CNN in previous applications, in this paper, we propose Xlnc1DCNN, a 1D-CNN model for classifying lncRNA and mRNA with explanation. The model solely uses nucleotide sequences as the training set. On the human test set, Xlnc1DCNN outperformed all other models in terms of accuracy and F1-Score. For the cross-species dataset, Xlnc1DCNN also had the generalization across testing species. We explained how the Xlnc1DCNN distinguished the lncRNA from mRNA transcript sequences by applying DeepSHAP to generate SHAP values representing what the model captured and visualized the contribution of each nucleotide using an in-house python code. The explanation of true positives (i.e., lncRNA transcript sequences) showed that the model classified a sequence as lncRNA if the sequence did not contain any important regions or contained only an N-terminal signal peptide or a single transmembrane helix. The explanation of true negatives (i.e., mRNA transcript sequences) showed that the model learned protein domains/families from the input transcript sequences and used them to predict the sequences as mRNAs. The explanation of false positives (i.e., mRNA predicted as lncRNA transcript sequences) showed that the model could not capture any important regions representing protein domains or families within these mRNA sequences. Some false positive sequences were also found with the inconsistent transcript types among the databases. Lastly, the explanation of false negatives (i.e., lncRNA predicted as mRNA transcript sequences) showed that the model captured protein domains or families within these lncRNA sequences, hence, misclassified them as mRNAs.

## 2. Results

### 2.1. Model Evaluation Results

We compared the performance of Xlnc1DCNN with eight existing tools: CPC2, CPAT, CNIT, PLEK, FEELnc, RNAsamba, LncADeep, and lncRNA_Mdeep [10–17] with the version listed in Supplementary Table S1. To have a fair and unbiased evaluation, we retrained CPAT, FEELnc, and RNAsamba that provided a training option using our human training dataset and used the pre-trained models of CPC2, CNIT, and LncADeep that did not provide a training option. Although PLEK and lncRNA_Mdeep came with a training option, retraining PLEK and lncRNA_Mdeep was very time-consuming, so we skipped to retrain both and used their default pre-trained models.

#### 2.1.1. Performance Evaluation on Human Test Set

The results on the human test set (Table 1) show that Xlnc1DCNN achieved the highest accuracy (94.53) and F1-Score (95.38), the second-highest precision (94.55) slightly lower than LncADeep, and the third-highest specificity (92.13) slightly lower than LncADeep and FEELnc. CPC2, CNIT, and CPAT achieved high sensitivity but much lower specificity. While FEELnc, RNAsamba, LncADeep, and lncRNA_Mdeep performed well on the average of every metric but overall, still lower than Xlnc1DCNN. We then analyzed the classification power of each tool by plotting a receiver operating characteristic curve (ROC) and measured area under the curve (AUC) as shown in Figure 1A, where Xlnc1DCNN achieved the highest AUC (0.9825) on the human test set. Figure 1B shows that Xlnc1DCNN also outperformed all tools on any range of sequence lengths of the human test set (Supplementary Table S2).

**Table 1.** Evaluation results of all tools on the human test set.

| Model | TP | FP | TN | FN | Accuracy | Sensitivity | Specificity | Precision | F1 |
| --- | --- | --- | --- | --- | --- | --- | --- | --- | --- |
| Xlnc1DCNN | 20,895 | 1,204 | 14,087 | 821 | **94.53** | 96.22 | 92.13 | 94.55 | **95.38** |
| CPC2 | 21,023 | 6,457 | 8,834 | 693 | 80.68 | 96.81 | 57.77 | 76.50 | 85.47 |
| CNIT | 21,307 | 3,580 | 11,711 | 409 | 89.22 | **98.12** | 76.59 | 85.61 | 91.44 |
| PLEK | 20,704 | 6,665 | 8,626 | 1,012 | 79.26 | 95.34 | 56.41 | 75.65 | 84.36 |
| CPAT | 20,646 | 2,597 | 12,694 | 1,070 | 90.09 | 95.07 | 83.02 | 88.83 | 91.84 |
| FEELNC | 20,023 | 1,182 | 14,109 | 1,693 | 92.23 | 92.20 | 92.27 | 94.43 | 93.30 |
| RNASAMBA | 20,998 | 1,795 | 13,496 | 718 | 93.21 | 96.69 | 88.26 | 92.12 | 94.35 |
| lncRNA_Mdeep | 20,813 | 1,799 | 13,492 | 903 | 92.70 | 95.84 | 88.23 | 92.04 | 93.90 |
| LncADeep | 20,232 | 1,113 | 14,178 | 1,484 | 92.98 | 93.17 | **92.72** | **94.79** | 93.97 |

**Figure 1:**
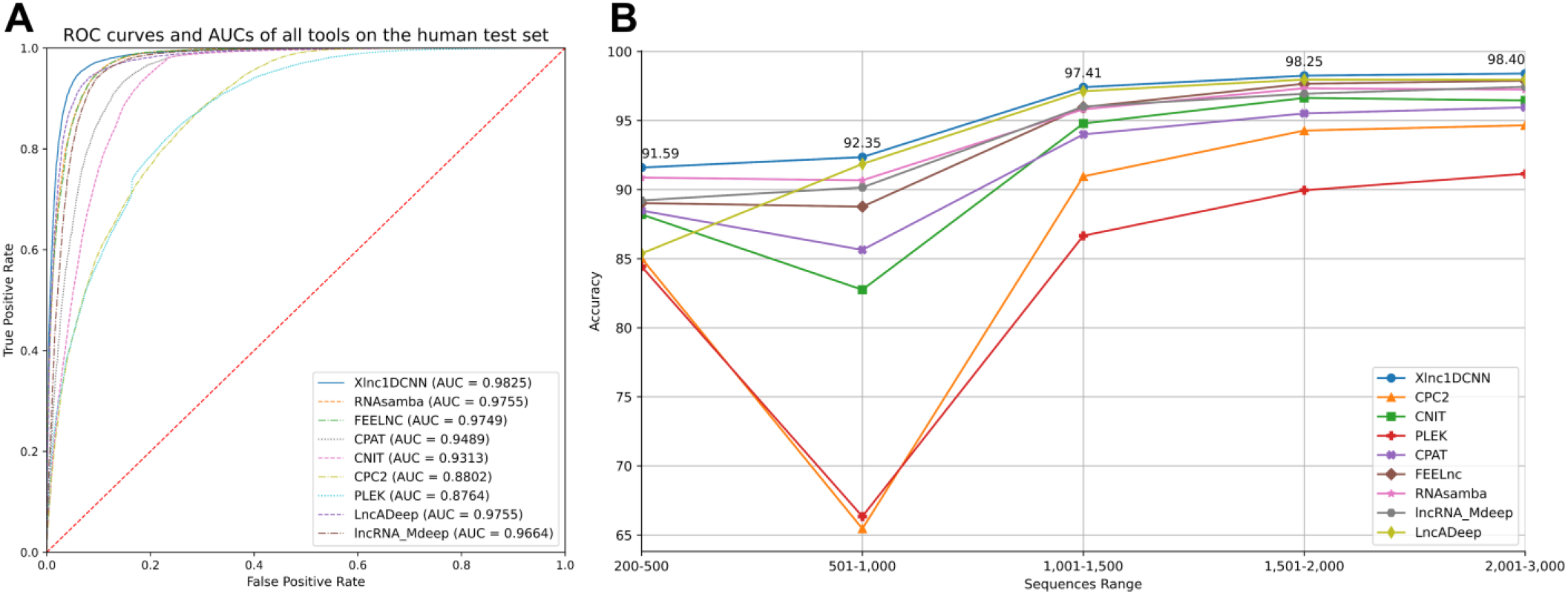
(**A**) ROC curve of all tools and their AUC on the human test set (**B**) Accuracy of all tools for any range of sequence lengths of the human test set.

#### 2.1.2. Performance Evaluation on Cross-Species Datasets

To evaluate the generalization of Xlnc1DCNN with cross-species datasets, we compared the model with other tools using mouse, gorilla, chicken, and cow datasets. The evaluation results show that Xlnc1DCNN, which was trained on the human dataset, has the generalization for classifying lncRNAs and mRNAs on other species (Table 2 and Supplementary Tables S3-S6). Xlnc1DCNN achieved the highest accuracy on gorilla dataset together with RNAsamba and the second highest accuracy on mouse dataset while LncADeep achieved the highest accuracy on mouse and cow datasets. Figure 2 shows that Xlnc1DCNN has the ROC curves and AUCs close to other tools on cross-species datasets. Overall, based on AUCs, LncADeep got the best generalization performance on cross-species datasets.

**Table 2.** Accuracy of nine models on cross-species datasets.

| Model | Mouse | Gorilla | Chicken | Cow |
| --- | --- | --- | --- | --- |
| Xlnc1DCNN | 92.58 | **96.06** | 92.35 | 95.92 |
| CPC2 | 80.06 | 94.96 | 93.51 | 94.48 |
| CNIT | 87.68 | 94.00 | 92.94 | 95.18 |
| PLEK | 73.62 | 89.53 | 79.54 | 86.22 |
| CPAT | 89.46 | 95.1 | 93.70 | 95.52 |
| FEELnc | 90.51 | 94.8 | 92.75 | 93.97 |
| RNASamba | 91.91 | **96.06** | **93.98** | 96.39 |
| LncADeep | **94.95** | 96.05 | 93.46 | **96.70** |
| lncRNA_Mdeep | 91.38 | 95.58 | 92.59 | 95.63 |

**Figure 2:**
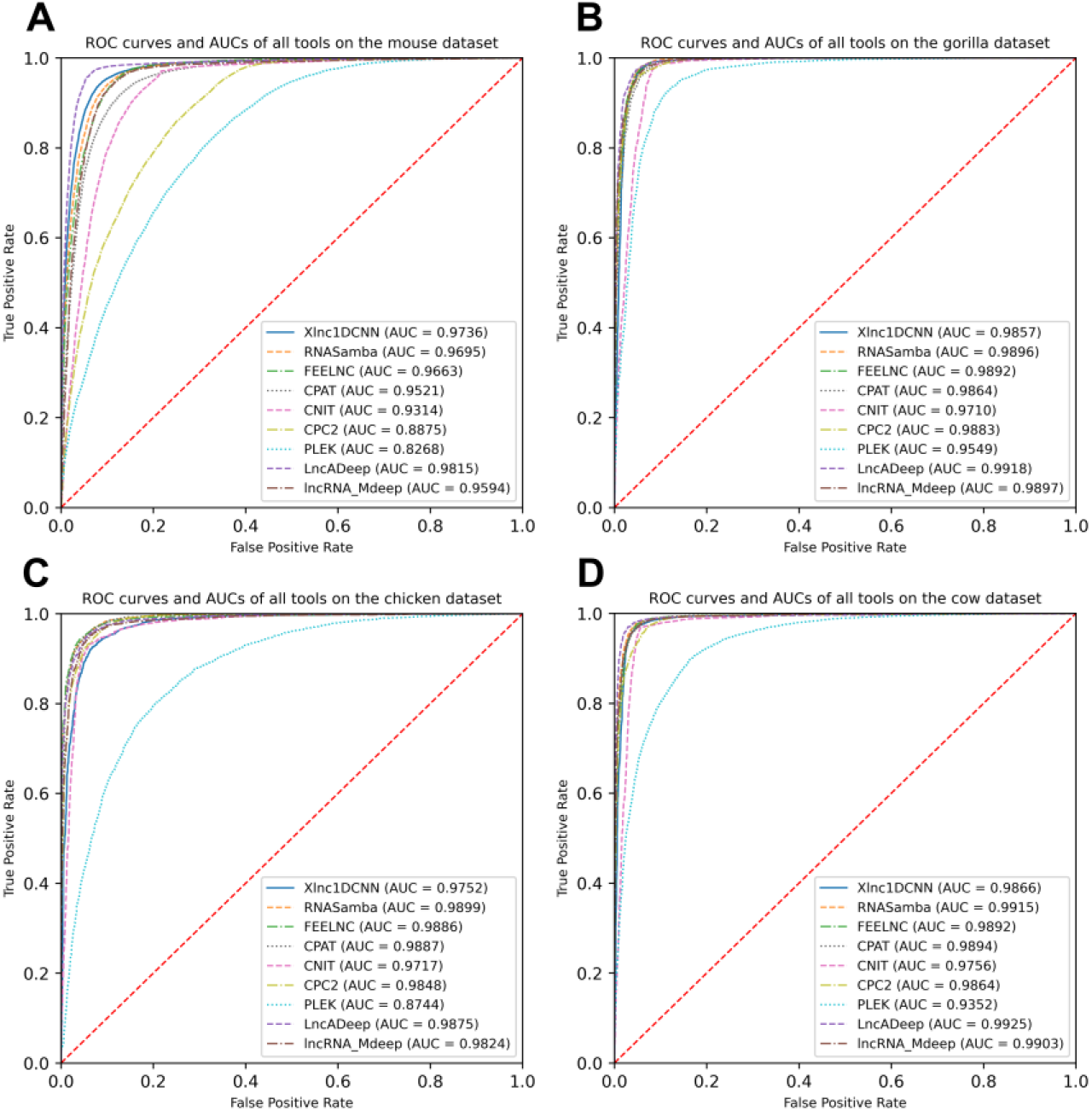
Receiver operating characteristic curves and AUCs of nine models on the datasets of (**A**) mouse (**B**) gorilla (**C**) cow and (**D**) chicken.

### 2.2. Model Interpretation Results

As Xlnc1DCNN outperformed other tools on the human test set, we assumed that 1D-CNN captured patterns within sequences that could be used to distinguish lncRNAs from mRNAs. To explain the model, we used DeepSHAP to describe the contribution of each nucleotide to the prediction results. The explanation output from DeepSHAP was SHAP values of all nucleotides of the entire sequence. This explanation result was then visualized based on the summed SHAP values of each three consecutive nucleotides, with important representative amino acids within the sequence highlighted.

In the following subsections, we present the explanation results of Xlnc1DCNN focusing on the true positive (TP), true negative (TN), false positive (FP), and false negative (FN) sequences predicted by Xlnc1DCNN on the human test set.

#### 2.2.1. True Positive Sequences

The explanation results of Xlnc1DCNN highlighted the important regions that contributed to the correct classification of an input lncRNA transcript sequence as a lncRNA as blue. From Figures 3A and 3B, the explanation results of the ENST00000658844.1 and lnc-REXO4-2:1 suggested that Xlnc1DCNN classified a transcript sequence as a lncRNA if it did not capture any important regions or specific patterns within the sequence. Additional explanation results of the TN sequences are shown in Supplementary Figures S1-S8.

**Figure 3.**
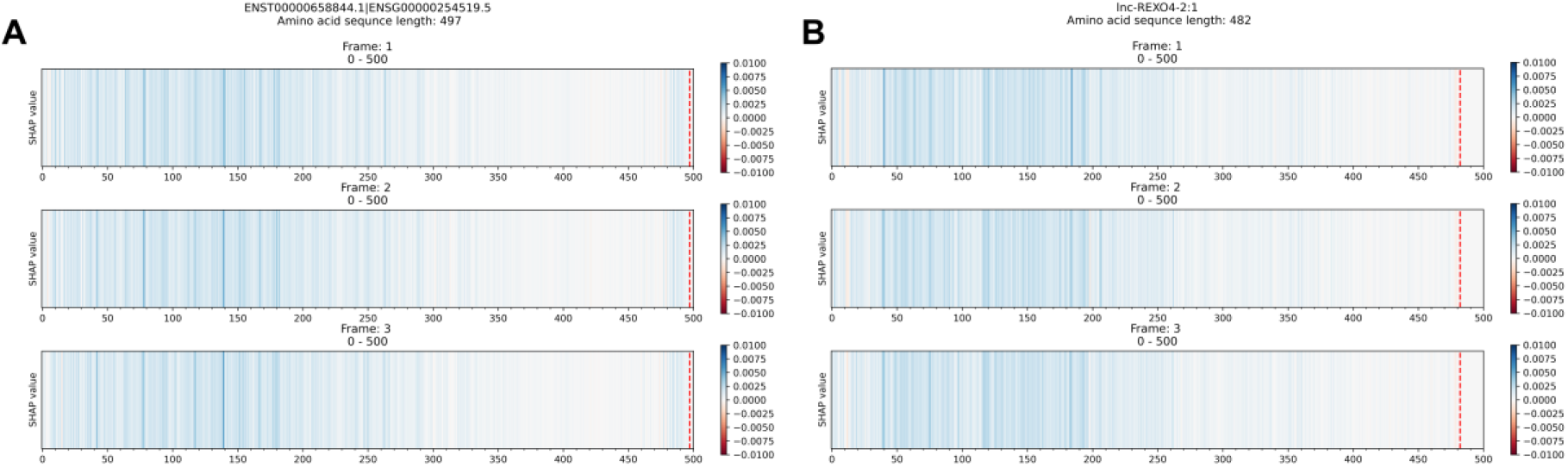
The explanation results of Xlnc1DCNN on TP sequences (A) ENST00000658844.1, a lncRNA sequence obtained from GENCODE and (B) lnc-REXO4-2:1, a lncRNA sequence obtained from LNCipedia.

#### 2.2.2. True Negative Sequences

The explanation results of Xlnc1DCNN highlighted the important regions of a protein-coding transcript (i.e., mRNA) as red as shown in Figures 4A-4C. To find out the importance of these regions, we utilized the available bioinformatics tools/databases such as TMHMM [28], Pfam [29], and InterPro [30] to identify protein domains or families in these sequences. Figure 4D shows the transmembrane helix regions of the ENST00000528724.5 transcript predicted by TMHMM, which correspond to the important regions captured by the Xlnc1DCNN. The prediction results of TMHMM and the explanation results of Xlnc1DCNN have similar patterns in several other mRNA transcripts within the test set (Supplementary Figures S9 and S10). Figure 4E shows the KRAB box (Krüppel associated box) identified by Pfam within the transcript ENST00000593088.5. This KRAB box is mostly overlapped with the important region captured by Xlnc1DCNN as shown in Figure 4B. Figure 4F shows the FAM32A family (family with sequence similarity 32 member A) identified by InterPro within the ENST00000589852.5 transcript. This corresponds to the important region of the ENST00000589852.5 identified by the Xlnc1DCNN as shown in Figure 4C. This transcript has been linked to ovarian tumor-associated gene [31]. Additional explanation results of the TN sequences are shown in Supplementary Figures S11-S14.

**Figure 4.**
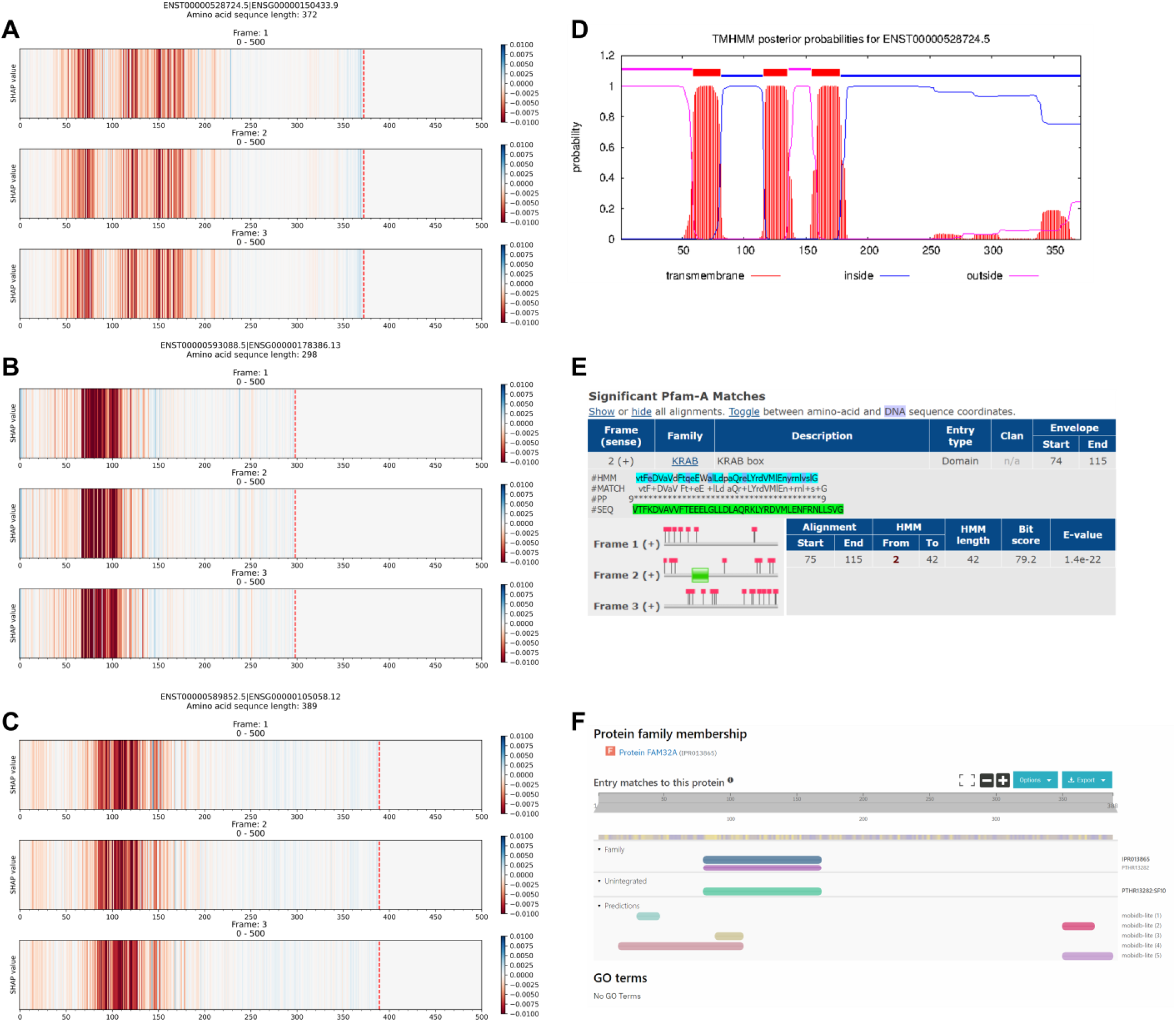
The comparison between the explanation results of Xlnc1DCNN on TN sequences (**A**) ENST00000528724.5, (**B**) ENST00000593088.5, and (**C**) ENST00000589852.5 protein-coding transcripts and (**D**) the prediction result of TMHMM program on the ENST00000528724.5 (**E**) the KRAB (Krüppel associated box) domain identified by Pfam within the ENST00000593088.5 and (**F**) the FAM32A family identified by InterPro within the ENST00000589852.5 transcripts.

#### 2.2.3. False Positive Sequences

False positive sequences are mRNA transcript sequences that are predicted as lncRNAs. Figure 5A shows the explanation result of ENST00000408930.6, which did not contain any important regions with red color contributing to the prediction as an mRNA. Figure 5B and 5C shows the result from Pfam and InterPro that both could not identify any protein domains or families within the ENST00000408930.6 protein-coding transcript also. While Ensembl database reports the ENST00000408930.6 as a protein-coding transcript of the HEPN1 (ENSG00000221932) gene, the Gene database at NCBI reports HEPN1 as the lncRNA gene (https://www.ncbi.nlm.nih.gov/gene/641654) and the RefSeq database reports the NR_170124.1 (ENST00000408930.6) as a long non-coding RNA (https://www.ncbi.nlm.nih.gov/nuccore/NR_170124.1). Based on our evaluation performance, the top five long non-coding identification tools (our Xlnc1DCNN, RNAsamba, LncADeep, lncRNA_Mdeep, FEELnc) predicted this sequence as lncRNA. This highlights an example of inconsistent annotations among public databases that affects the model performance and evaluation. Additional explanation results of the FP sequences are shown in Supplementary Figures S15-S19.

**Figure 5.**
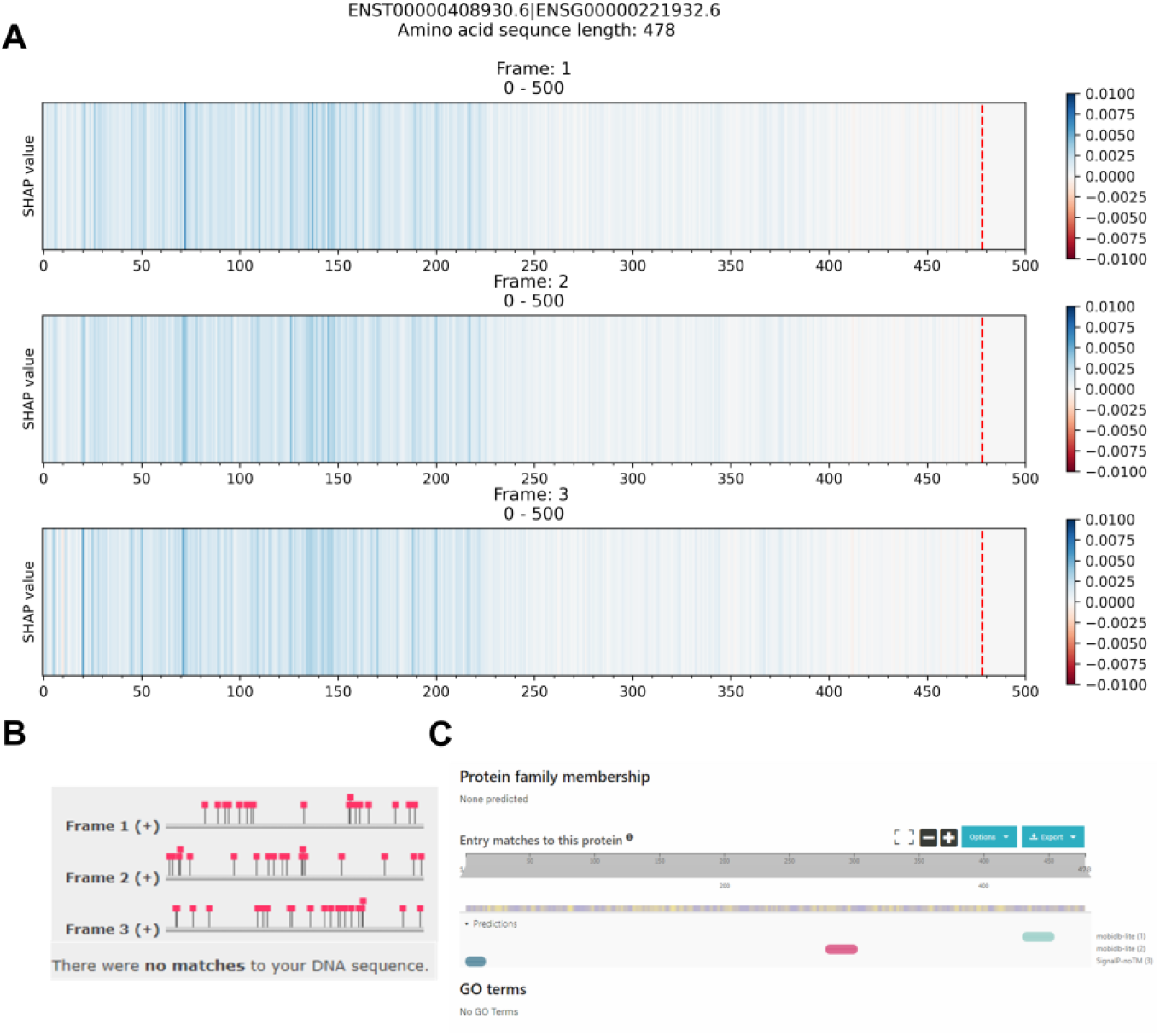
The comparison between (**A**) the explanation result of Xlnc1DCNN on the ENST00000408930.6 protein-coding transcript, predicted as a lncRNA (**B**) the identification result from Pfam and (**C**) the identification result from InterPro.

#### 2.2.4. False Negative Sequences

False negative sequences are lncRNA transcript sequences that are predicted as mRNAs. Figure 6A and 6B shows the explanation results of lncRNAs: LNC-SIGIRR-2:1 and ENST00000616537.4 with important regions that contributed to the wrong prediction as mRNA transcripts. These regions correspond to the identified Anoctamin and the Taxilin InterPro families identified by InterPro as shown in Figures 6C and 6D. Additional explanation results of the FN sequences are shown in Supplementary Figures S20-S25.

**Figure 6.**
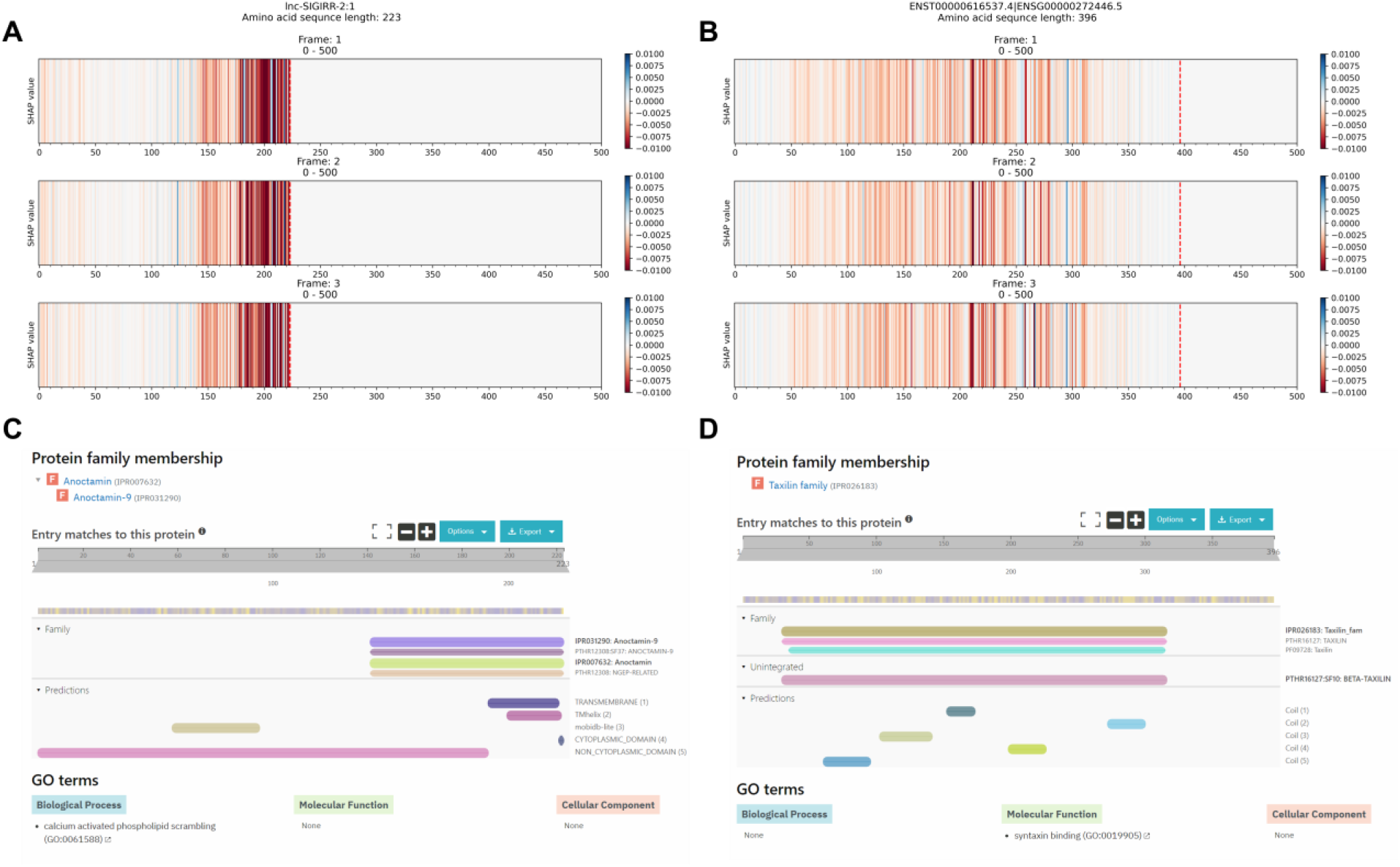
The comparison between the explanation result of Xlnc1DCNN on the long non-coding RNA transcripts (**A**) lnc-SIGIRR-2:1 and (**B**) ENST00000616537.4, predicted as mRNAs (**C**) the Anoctamin family within the lnc-SIGIRR-2:1 transcript and (**D**) the Taxilin family within the ENST00000616537.4 transcript identified by InterPro.

## 3. Discussion

The explanation results of Xlnc1DCNN on the true positive sequences show that most of the lncRNAs were found with no conserved regions or patterns in short regions with unknown functions (Supplementary Figures S1, S2). The important regions of some other lncRNA sequences highlighted a transmembrane helix (Supplementary Figures S3-S5) or signal peptide (Supplementary Figures S6-S8). Over recent years, some studies also found transmembrane helix inside lncRNAs [32,33] and hidden peptides encoded within non-coding RNAs [34]. These findings correspond to what Xlnc1DCNN has learned and highlighted via the explanation result as important regions for classifying a sequence as lncRNA.

On the true negative sequences, the explanation results of Xlnc1DCNN show that the model could capture the regions representing the protein domains or families in the transcript sequences. Hence, it could classify most of the input mRNA sequences correctly as the protein-coding transcripts.

Most of the explanation results of false positive sequences do not contain the important regions (red color) that contributed to the model prediction as mRNAs. Pfam and InterPro also could not identify any protein domains and/or families within these sequences. Some of the false positive sequences (e.g., ENST00000567119.1) were found with only intrinsically disordered regions (IDRs). In addition, we also found that the biotype, classified by Ensembl, of several false positive sequences (e.g., ENST00000637131.1, ENST00000540988.1) are categorized as nonsense-mediated decay or processed transcript (i.e., gene/transcript that does not contain an open reading frame), and some sequences (e.g., ENST00000408930.6) have much shorter length of protein sequence compared to their transcript sequence.

Most of the explanation results of false negative sequences do contain important regions that correspond to Pfam or InterPro domains. Supplementary Figures S20-S25 show additional examples of lncRNA sequences that were misclassified as mRNAs by all top five tools.

As recent studies found that some putative lncRNAs contain a short open reading frame (sORF) [35], we further analyzed the association of lncRNAs and sORF using the explanation results of Xlnc1DCNN. Some false negative sequences were randomly selected and checked if they contained sORF by using MetamORF [36]. While MetamORF found sORFs in some of these sequences, the reported regions of these sORFs did not correspond to the important regions highlighted by the explanation results.

## 4. Materials and Methods

### 4.1 Data Compilation and Pre-processing

The human transcript datasets for training the model were obtained from GENCODE [37] and LNCipedia [38]. GENCODE (release 32) contains 48,351 sequences of lncRNA transcripts and 100,291 sequences of protein-coding transcripts (PCTs). For LNCipedia (version 5.2), only high confidence sequences were selected, which resulted in 107,039 lncRNA transcripts. To remove lncRNA transcript sequences from LNCipedia that are duplicates of GENCODE, we used CD-HIT-EST-2D [39] to compare lncRNA sequences between LNCipedia and GENCODE and filtered out the sequences with more than 95% similarity from LNCipedia dataset. A total of 72,803 lncRNA sequences from LNCipedia remained. We then pre-processed the sequences used for training the Xlnc1DCNN model by discarding the sequences shorter than 200 bases and longer than 3,000 bases. After filtering, one-hot encoding was used to encode the sequences. The total number of remaining sequences after cleansing was 185,030 with the 108,578 lncRNAs and 76,453 PCTs (Table 3). The lncRNAs and PCTS were set as the positive and negative class, respectively. The dataset was stratified split by 80% and 20 % into the training and test sets.

**Table 3.** The summary of datasets from GENCODE and LNCipedia.

| Sequence | Species | Data Source | Dataset Size | < 200 bps | > 3,000 bps | Number of transcripts after cleansing |
| --- | --- | --- | --- | --- | --- | --- |
| mRNA | Human | GENCODE (release 32) | 100,291 | 374 | 23,464 | 76,453 |
| lncRNA | Human | GENCODE (release 32) | 48,351 | 291 | 3,486 | 44,574 |
| lncRNA | Human | LNCipedia (version 5.2) | 72,803 | 0 | 8,799 | 64,004 |

Cross-species datasets included the mouse dataset obtained from GENCODE [37] release (M23) and the gorilla, chicken, and cow datasets obtained from Ensembl [40] (release 102). We pre-processed the cross-species datasets by discarding the sequences shorter than 200 bases and longer than 3,000 bases. We then randomly selected the mRNA and lncRNA sequences for each species. The test transcripts of gorilla, chicken, cow, and mouse contained 8,000, 8,000, 11,000, and 32,000 sequences, each with the equal number of sequences from each class.

### 4.2. Model Architecture

In this study, we designed and implemented the Xlnc1DCNN model in Python3 using TensorFlow on NVIDIA GeForce GTX 1080 Ti and Intel Xeon Silver 4112 Processor. The built model could distinguish lncRNAs from the mRNAs (PCTs) and outperformed the existing tools for the human dataset. The model architecture consists of three convolutions with pooling layers, two fully connected layers, and a Softmax layer. We used ReLU as the activation function for convolution and fully connected layers. We also found that adding the dropout layer after the pooling layer made the model perform slightly better.

#### 4.2.1. Hyperparameter Optimization and Details of Learning

We used 10% of the data from the training set to perform hyperparameter optimizations over the kernel size, dropout rate, stride size, batch size, and learning rate by using the grid search algorithm. The best kernel size was 57, with the stride size equal to 1. The model performance started to decrease after increasing the stride size for almost every kernel size. For the learning details, the momentum, learning rate, number of epochs, and batch size were 0.9, 0.01, 120, and 128, with the stochastic gradient descend as an optimizer. The final hypermeters used in the model architecture are shown in Table 4.

**Table 4.** The hyperparameters of proposed 1D-CNN architecture.

| Layer | Hyperparameter |
| --- | --- |
| Conv 1D | kernel size=57, stride=1 |
| Max-Pooling | pool size=2 |
| Dropout | p=0.3 |
| Conv 1D | kernel size=57, stride=1 |
| Max-Pooling | pool size=2 |
| Dropout | p=0.3 |
| Conv 1D | kernel size=57, stride=1 |
| Max-Pooling | pool size=2 |
| Dropout | p=0.3 |
| Flatten | - |
| Dense | 256 |
| Dropout | p=0.5 |
| Dense | 256 |
| Dropout | p=0.5 |
| Softmax | 2 |

### 4.3 Model Interpretation

DeepSHAP was used to interpret how the proposed Xlnc1DCNN model could classify the lncRNAs and mRNAs from the input transcript sequences. As DeepSHAP needs background distributions as references to approximate the SHAPley values on conditional expectation, 175 sequences from each class were randomly selected as the representative background. The total 350 sequences were used as the backgrounds as it was limited by the available GPU.

The output from DeepSHAP is SHAP values which represent the contribution of each nucleotide from the model. To obtain SHAP values representing each nucleotide within a sequence, we summed up SHAP values inside the array of one-hot encoding and got a single SHAP value of each nucleotide. To visualize SHAP values from DeepSHAP of the input transcript sequence, we further summed up the SHAP values of three consecutive nucleotides, which probably represented an amino acid, and generated the results in three reading frames. We then plotted a color line for each representative amino acid. The blue and red color respectively indicates the contribution of each amino acid to the classification of a sequence as a lncRNA and an mRNA (Figure 7).

**Figure 7.**
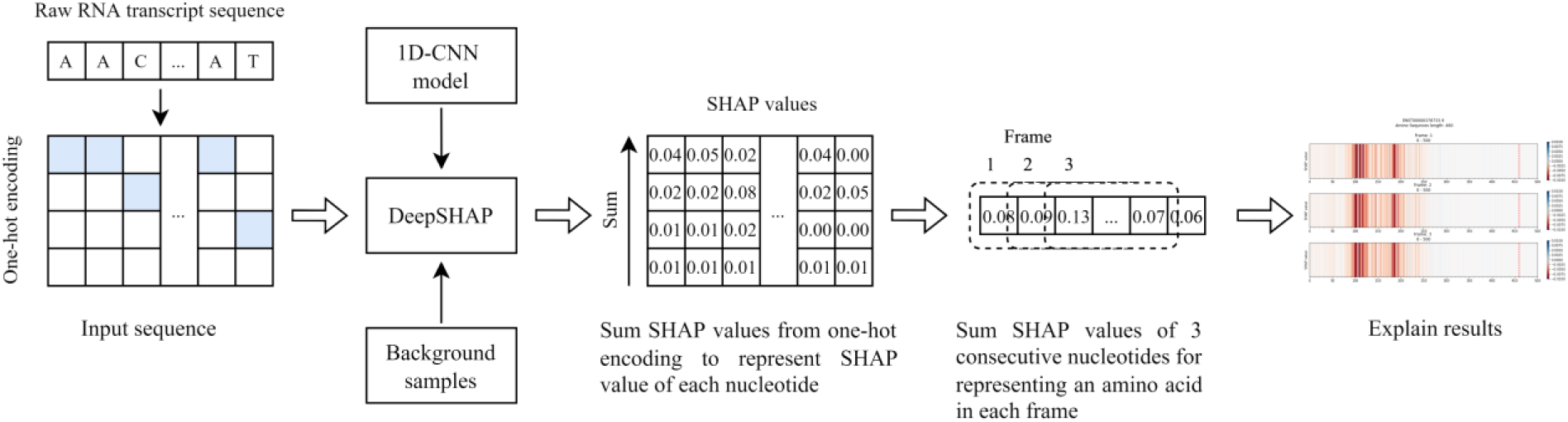
The process to obtain SHAP values for explaining the nucleotide contribution that was captured by the model to differentiate lncRNA from mRNA transcript sequences.

### 4.4 Evaluation Metrics

To evaluate the performance of the proposed Xlnc1DCNN model with other existing tools, we used the following metrics. TP are the lncRNA transcript sequences that are predicted as lncRNAs, and TN are PCTs that are predicted as PCTs. FP are the PCTs that are predicted as lncRNAs while FN are lncRNAs that are predicted as PCTs.

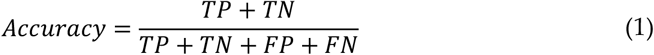

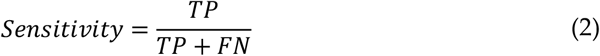

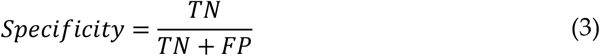

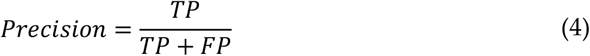

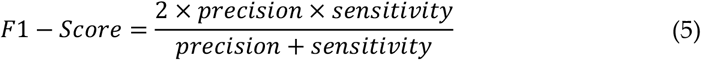

## 5. Conclusions

In this study, we proposed Xlnc1DCNN, a simple but effective 1D-CNN model for classification and explanation of lncRNA and protein-coding transcripts. We have shown that using 1D-CNN as a feature extractor can lead to a better prediction performance than other existing tools using traditional feature extraction methods. The explanation results provided several insights of what the model learned to distinguish the lncRNA from protein-coding transcripts. A single transmembrane helix region highlighted by the explanation results of several true positive lncRNA transcripts agreed with the recent findings of transmembrane microproteins within lncRNAs. Disordered proteins without any important regions highlighted in the explanation results were misclassified as lncRNAs. Several explanation results of lncRNA misclassified as protein-coding transcripts contained important regions that correspond to protein domains or families in Pfam and/or InterPro. These insights revealed the complexity in long non-coding RNAs and the need of periodic evaluation of cross-referenced gene annotation among public databases.

