## Supplementary for "Interpretable Deep Learning Model Reveals Subsequences of Various Functions for Long Non-coding RNA Identification"

**Table S1.** Software version.

| Software | Version |
|---|---|
| CPC2 | Web based |
| CNIT | Web based |
| PLEK | 1.2 |
| CPAT | 2.0.0 |
| FEELnc | 0.1.1 |
| RNAsamba | 0.2.4 |
| LncADeep | 1.0 |
| lncRNA_Mdeep | - |

**Table S2.** The number of mRNAs and lncRNAs of each range of sequences on the human test set.

| Sequence Range | Number of mRNAs | Number of lncRNAs |
|---|---|---|
| 200–500 | 1,149 | 6,049 |
| 501–1,000 | 6,666 | 7,914 |
| 1,001–1,500 | 2,124 | 3,438 |
| 1,501–2,000 | 2,189 | 2,090 |
| 2,000–3,000 | 3,163 | 2,225 |

**Table S3.** Evaluation results of all tools on gorilla transcripts.

| Model | TP | FP | TN | FN | Accuracy | Sensitivity | Specificity | Precision | F1-Score |
|---|---|---|---|---|---|---|---|---|---|
| Xlnc1DCNN | 3,902 | 217 | 3,783 | 98 | 96.06 | 97.55 | 94.58 | 94.73 | 96.12 |
| CPC2 | 3,878 | 281 | 3,719 | 122 | 94.96 | 96.95 | 92.98 | 93.24 | 95.06 |
| CNIT | 3,944 | 424 | 3,576 | 56 | 94.00 | 98.60 | 89.40 | 90.29 | 94.26 |
| PLEK | 3,847 | 685 | 3,315 | 153 | 89.53 | 96.18 | 82.88 | 84.89 | 90.18 |
| CPAT | 3,824 | 216 | 3,784 | 176 | 95.10 | 95.60 | 94.60 | 94.65 | 95.12 |
| FEELnc | 3,723 | 139 | 3,861 | 277 | 94.80 | 93.08 | 96.53 | 96.40 | 94.71 |
| RNAsamba | 3,899 | 214 | 3,786 | 101 | 96.06 | 97.48 | 94.65 | 94.80 | 96.12 |
| lncRNA_Mdeep | 3,863 | 217 | 3,783 | 137 | 95.58 | 96.58 | 94.58 | 94.68 | 95.62 |
| LncADeep | 3,842 | 158 | 3,842 | 158 | 96.05 | 96.05 | 96.05 | 96.05 | 96.05 |

**Table S4.** Evaluation results of all tools on chicken transcripts.

| Model | TP | FP | TN | FN | Accuracy | Sensitivity | Specificity | Precision | F1-Score |
|---|---|---|---|---|---|---|---|---|---|
| Xlnc1DCNN | 3,606 | 218 | 3,782 | 394 | 92.35 | 90.15 | 94.55 | 94.30 | 92.18 |
| CPC2 | 3,679 | 198 | 3,802 | 321 | 93.51 | 91.98 | 95.05 | 94.89 | 93.41 |
| CNIT | 3,737 | 302 | 3,698 | 263 | 92.94 | 93.43 | 92.45 | 92.52 | 92.97 |
| PLEK | 3,119 | 756 | 3,244 | 881 | 79.54 | 77.98 | 81.10 | 80.49 | 79.21 |
| CPAT | 3,593 | 97 | 3,903 | 407 | 93.70 | 89.83 | 97.58 | 97.37 | 93.45 |
| FEELnc | 3,485 | 65 | 3,935 | 515 | 92.75 | 87.13 | 98.38 | 98.17 | 92.32 |
| RNAsamba | 3,618 | 100 | 3,900 | 382 | 93.98 | 90.45 | 97.50 | 97.31 | 93.75 |
| lncRNA_Mdeep | 3,538 | 131 | 3,869 | 462 | 92.59 | 88.45 | 96.73 | 96.43 | 92.27 |
| LncADeep | 3,586 | 109 | 3,891 | 414 | 93.46 | 89.65 | 97.28 | 97.05 | 93.20 |

**Table S5.** Evaluation results of all tools on mouse transcripts.

| Model | TP | FP | TN | FN | Accuracy | Sensitivity | Specificity | Precision | F1 |
|---|---|---|---|---|---|---|---|---|---|
| Xlnc1DCNN | 15,307 | 1,680 | 14,320 | 693 | 92.58 | 95.67 | 89.50 | 90.11 | 92.81 |
| CPC2 | 15,186 | 5,568 | 10,432 | 814 | 80.06 | 94.91 | 65.20 | 73.17 | 82.64 |
| CNIT | 15,530 | 3,473 | 12,527 | 470 | 87.68 | 97.06 | 78.29 | 81.72 | 88.74 |
| PLEK | 14,731 | 7,172 | 8,828 | 1,269 | 73.62 | 92.07 | 55.18 | 67.26 | 77.73 |
| CPAT | 14,812 | 2,186 | 13,814 | 1,188 | 89.46 | 92.58 | 86.34 | 87.14 | 89.78 |
| FEELnc | 14,244 | 1,281 | 14,719 | 1,756 | 90.51 | 89.03 | 91.99 | 91.75 | 90.37 |
| RNAsamba | 15,161 | 1,749 | 14,251 | 839 | 91.91 | 94.76 | 89.07 | 89.66 | 92.14 |
| lncRNA_Mdeep | 14,858 | 1,616 | 14,384 | 1,142 | 91.38 | 92.86 | 89.90 | 90.19 | 91.51 |
| LncADeep | 15,388 | 1,003 | 14,997 | 612 | 94.95 | 96.18 | 93.73 | 93.88 | 95.01 |

**Table S6.** Evaluation results of all tools on cow transcripts.

| Model | TP | FP | TN | FN | Accuracy | Sensitivity | Specificity | Precision | F1 |
|---|---|---|---|---|---|---|---|---|---|
| Xlnc1DCNN | 5,259 | 208 | 5,292 | 241 | 95.92 | 95.62 | 96.22 | 96.20 | 95.91 |
| CPC2 | 5,201 | 308 | 5,192 | 299 | 94.48 | 94.56 | 94.40 | 94.41 | 94.49 |
| CNIT | 5,324 | 354 | 5,146 | 176 | 95.18 | 96.80 | 93.56 | 93.77 | 95.26 |
| PLEK | 4,751 | 767 | 4,733 | 749 | 86.22 | 86.38 | 86.05 | 86.10 | 86.24 |
| CPAT | 5,194 | 187 | 5,313 | 306 | 95.52 | 94.44 | 96.60 | 96.52 | 95.47 |
| FEELnc | 4,491 | 124 | 5,376 | 509 | 93.97 | 89.82 | 97.75 | 97.31 | 93.42 |
| RNAsamba | 5,294 | 191 | 5,309 | 206 | 96.39 | 96.25 | 96.53 | 96.52 | 96.39 |
| lncRNA_Mdeep | 5,188 | 169 | 5,331 | 312 | 95.63 | 94.33 | 96.93 | 96.85 | 95.57 |
| LncADeep | 5,262 | 121 | 5,379 | 238 | 96.74 | 95.67 | 97.80 | 97.75 | 96.70 |

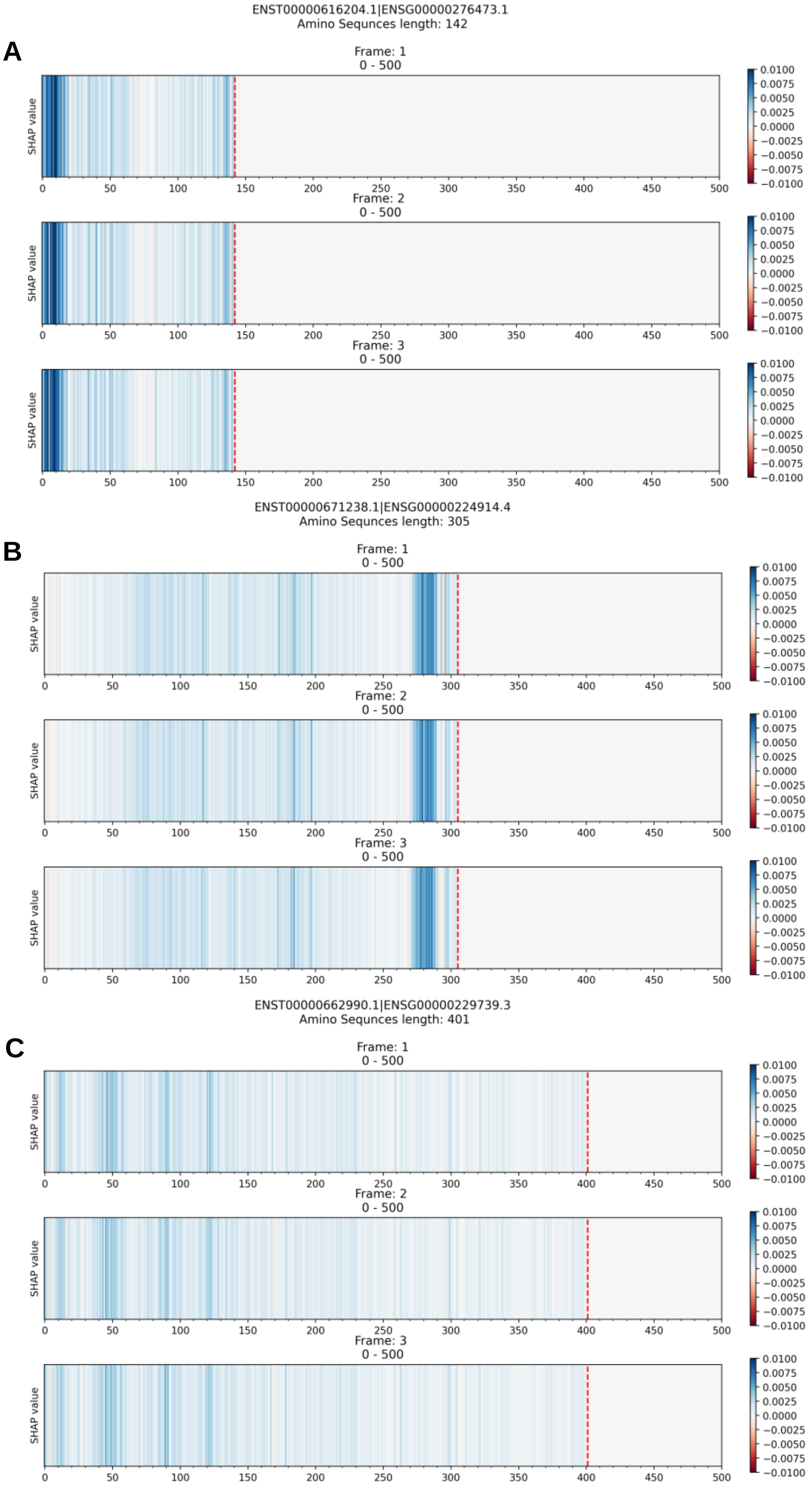


**Figure S1.** The explanation results of Xlnc1DCNN on true positive sequences of the (A) ENST00000616204.1, (B) ENST00000671238.1, and (C) ENST00000662990.1 that obtained from GENCODE.

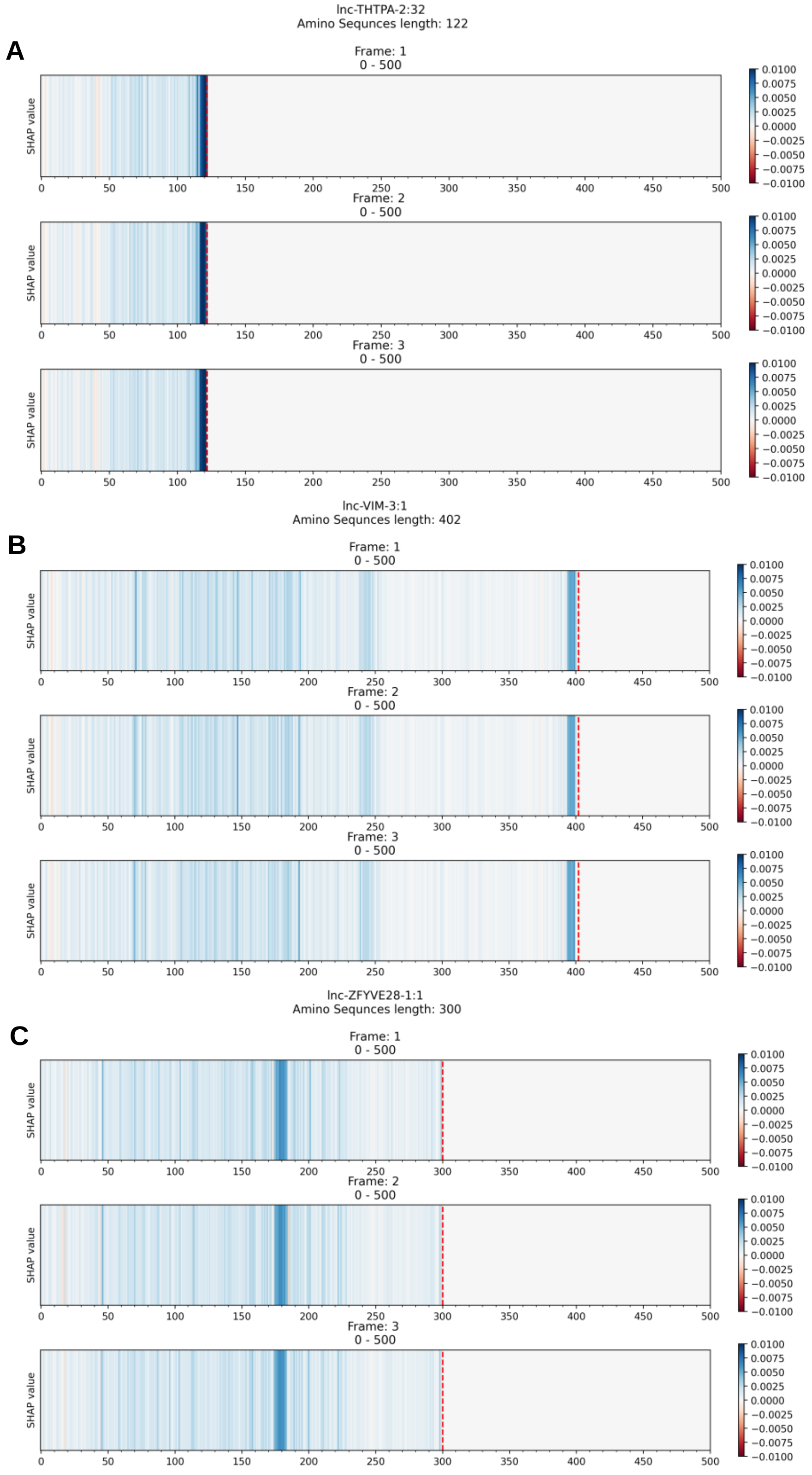


**Figure S2.** The explanation results of Xlnc1DCNN on true positive sequences of the (**A**) lnc-THTPA-2:32, (**B**) lnc-VIM-3:1, and (**C**) lnc-ZFYVE28-1:1 that obtained from LNCipedia.

A

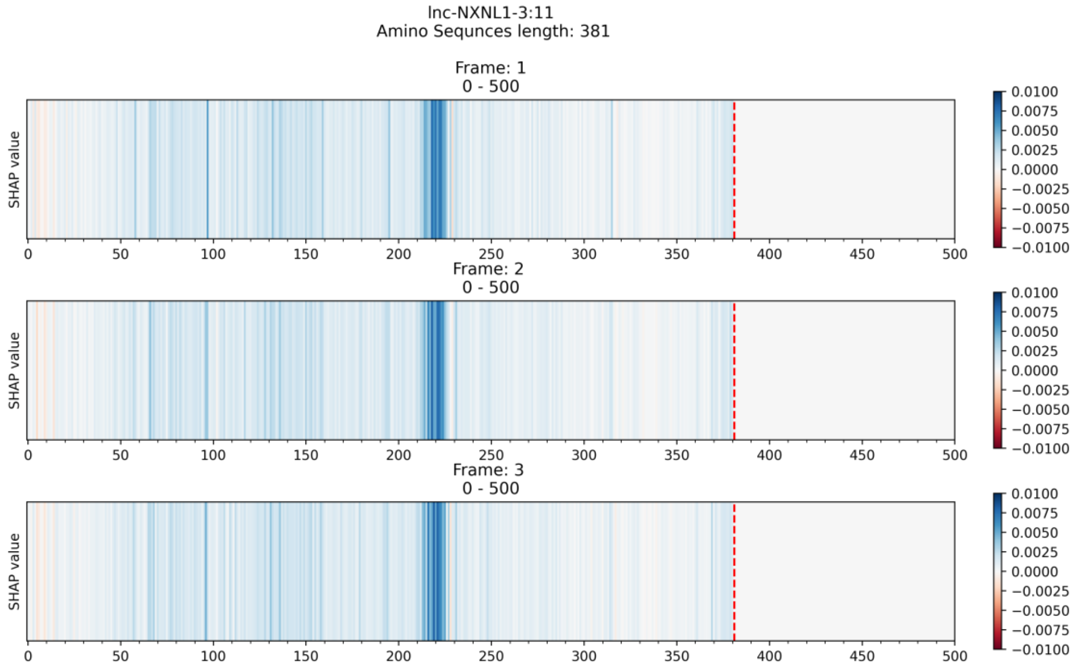


B

```
# lnc-NXNL1-3:11 Length: 381
# lnc-NXNL1-3:11 Number of predicted TMHs:  1
# lnc-NXNL1-3:11 Exp number of AAs in TMHs: 19.88156
# lnc-NXNL1-3:11 Exp number, first 60 AAs:  0
# lnc-NXNL1-3:11 Total prob of N-in:        0.93032
lnc-NXNL1-3:11  TMHMM2.0        inside       1   214
lnc-NXNL1-3:11  TMHMM2.0        TMhelix    215   232
lnc-NXNL1-3:11  TMHMM2.0        outside    233   381
```

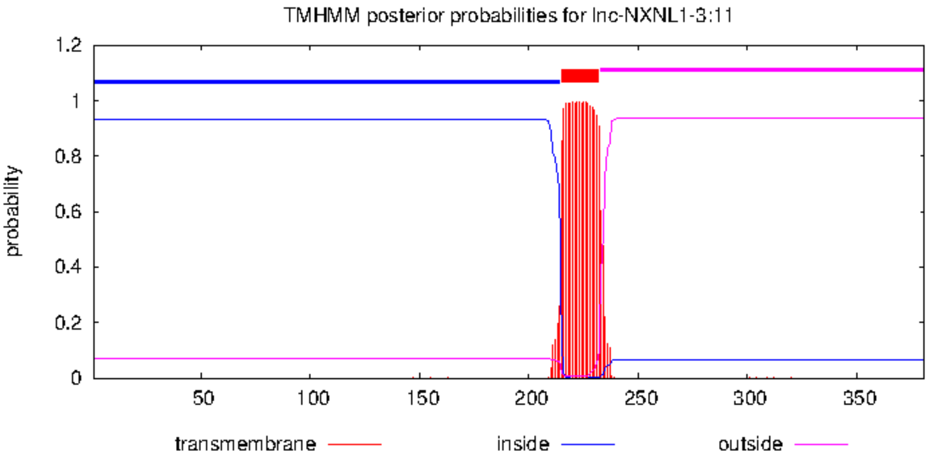


**Figure S3.** (A) The explanation result of Xlnc1DCNN and (B) the prediction result of the transmembrane helices by TMHMM program, on the true positive sequence, lnc-NXNL1-3:11.

**A**

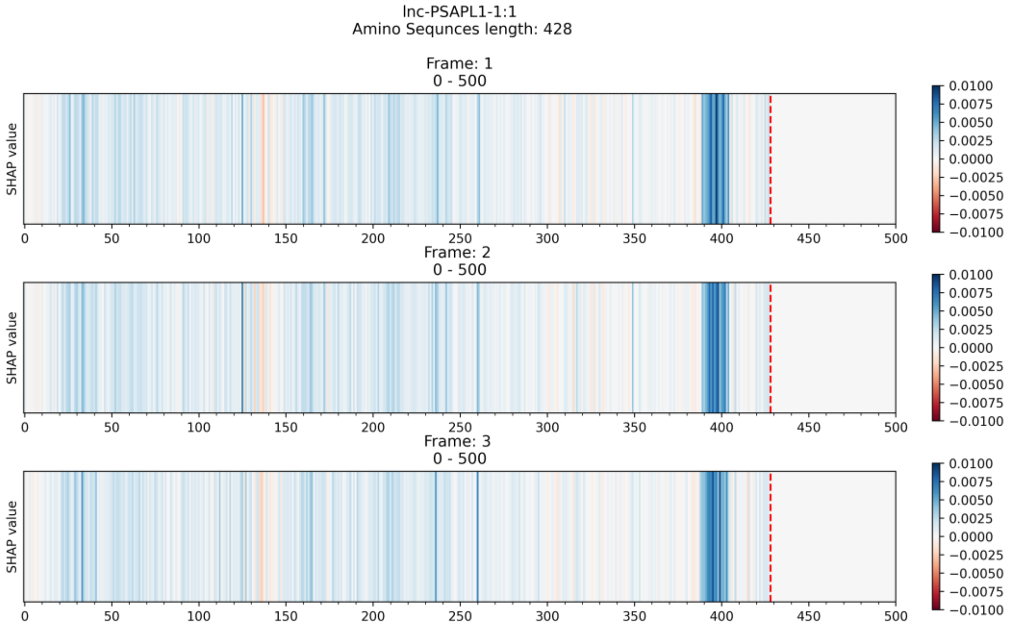


**B**

```
# lnc-PSAPL1-1:1 Length: 428
# lnc-PSAPL1-1:1 Number of predicted TMHs:  1
# lnc-PSAPL1-1:1 Exp number of AAs in TMHs: 23.24178
# lnc-PSAPL1-1:1 Exp number, first 60 AAs:  0
# lnc-PSAPL1-1:1 Total prob of N-in:        0.00991
lnc-PSAPL1-1:1  TMHMM2.0        outside        1   388
lnc-PSAPL1-1:1  TMHMM2.0        TMhelix      389   411
lnc-PSAPL1-1:1  TMHMM2.0        inside       412   428
```

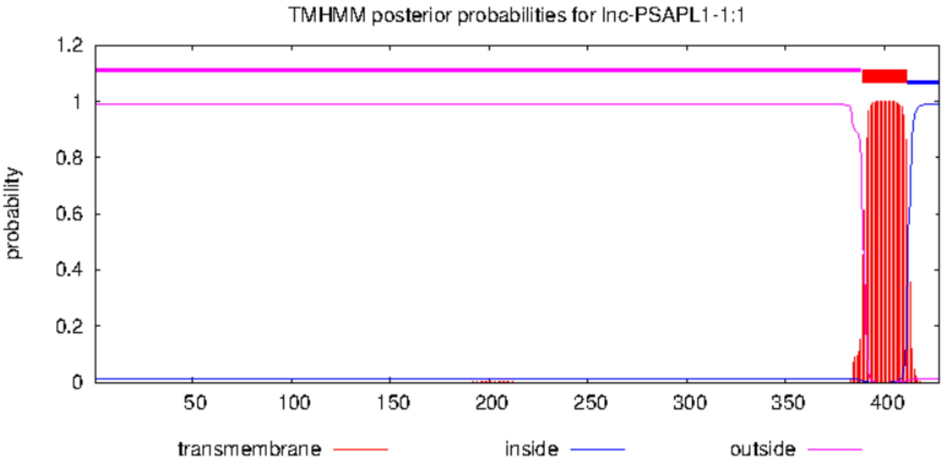


**Figure S4.** (**A**) The explanation result of Xlnc1DCNN and (**B**) the prediction result of the transmembrane helices by TMHMM program, on the true positive sequence, lnc-PSAPL1-1:1.

**A**

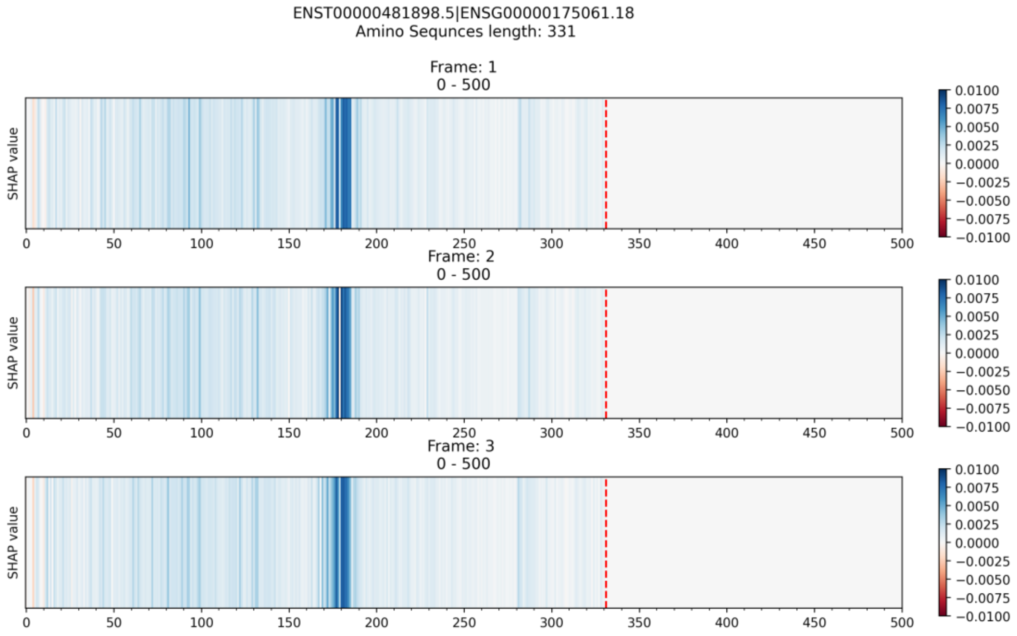


**B**

```
# ENST00000481898.5 Length: 330
# ENST00000481898.5 Number of predicted TMHs:  1
# ENST00000481898.5 Exp number of AAs in TMHs: 25.62541
# ENST00000481898.5 Exp number, first 60 AAs:  0
# ENST00000481898.5 Total prob of N-in:        0.04000
ENST00000481898.5   TMHMM2.0    outside       1   166
ENST00000481898.5   TMHMM2.0    TMhelix     167   186
ENST00000481898.5   TMHMM2.0    inside      187   330
```

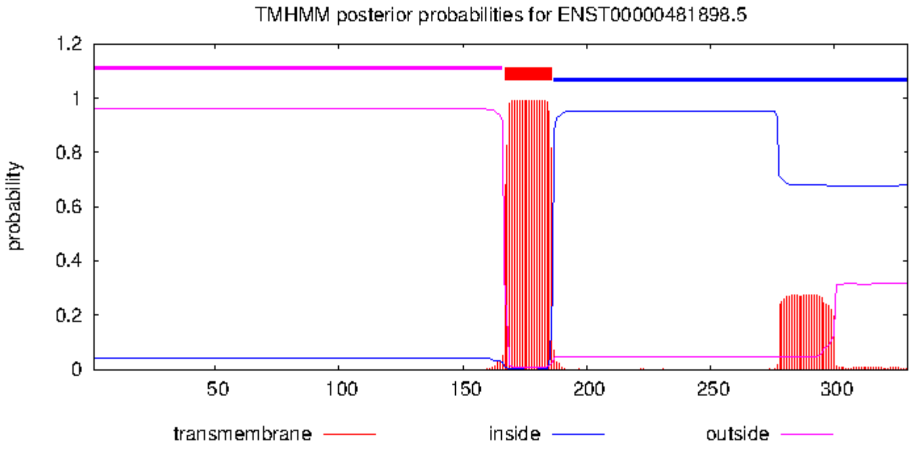


**Figure S5.** (**A**) The explanation result of Xlnc1DCNN and (**B**) the prediction result of the transmembrane helices by TMHMM program, on the true positive sequence, ENST00000481898.5.

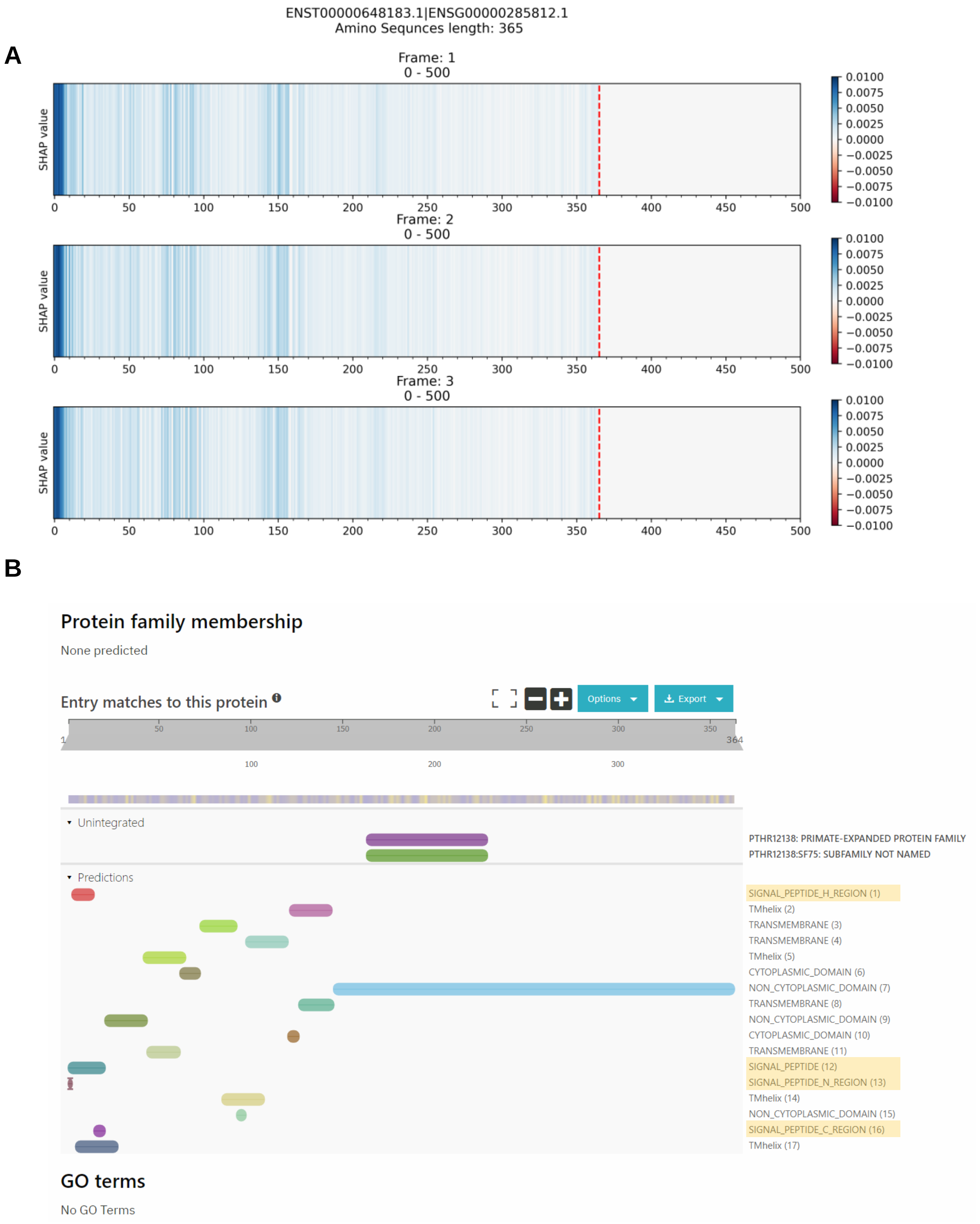


**Figure S6.** (**A**) The explanation result of Xlnc1DCNN and (**B**) the signal peptide identified by InterPro, on the true positive sequence, ENST00000648183.1.

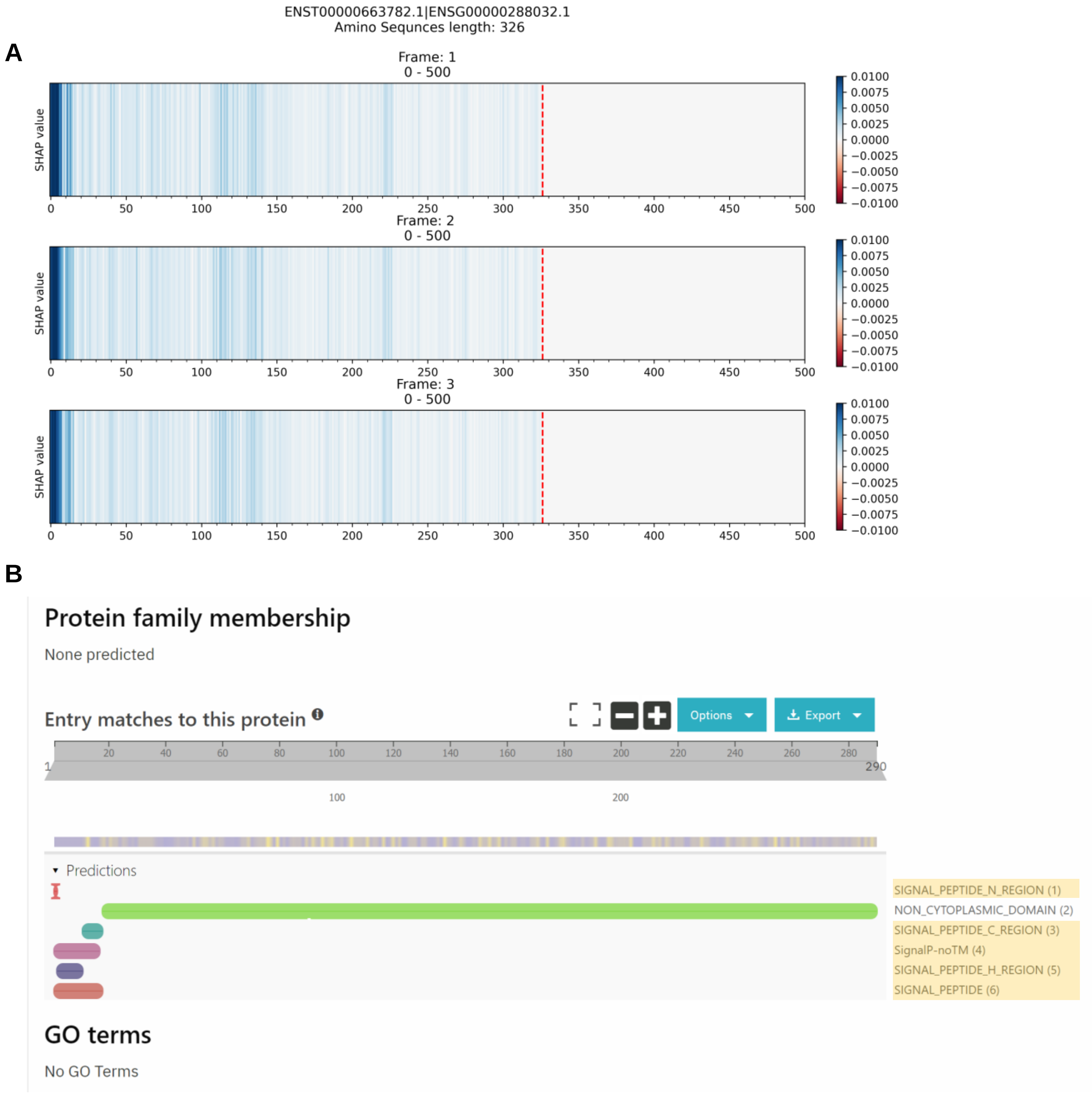


**Figure S7.** (A) The explanation result of Xlnc1DCNN and (B) the signal peptide identified by InterPro, on the true positive sequence, ENST00000663782.1.

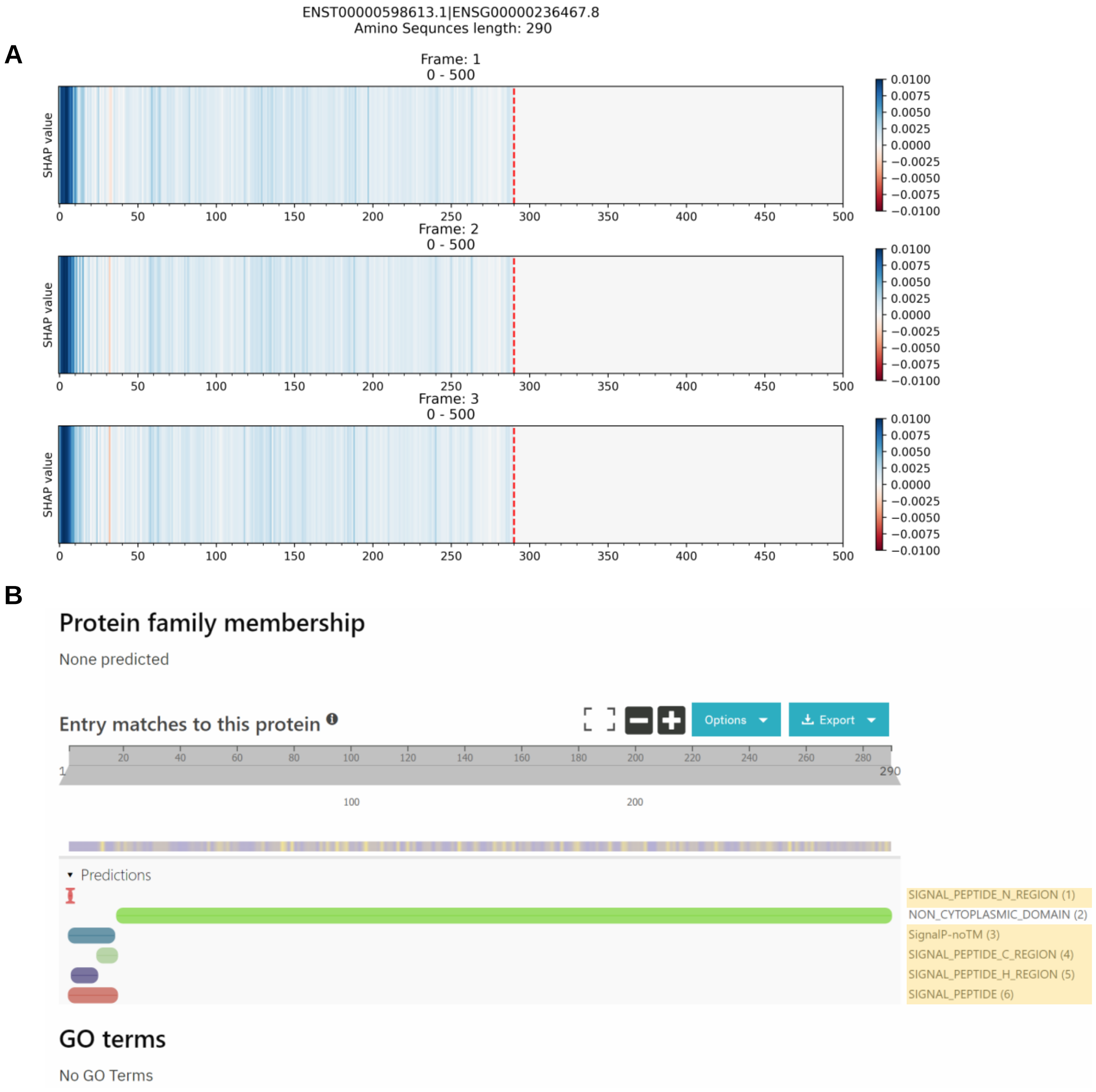


**Figure S8.** (A) The explanation result of Xlnc1DCNN and (B) the signal peptide identified by InterPro, on the true positive sequence, ENST00000598613.1.

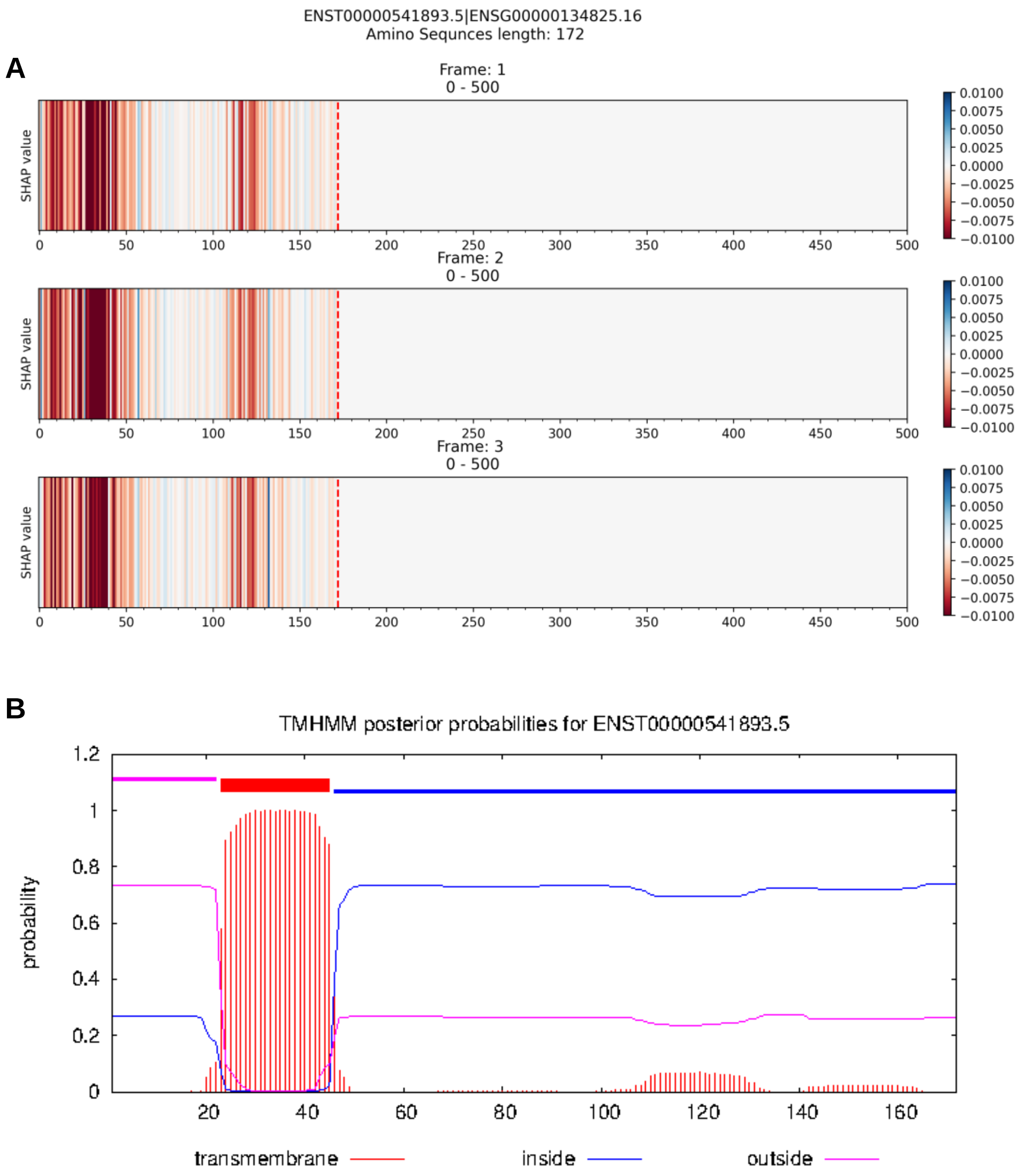


**Figure S9.** (A) The explanation result of Xlnc1DCNN and (B) the prediction result of the transmembrane helices by TMHMM program, on the true negative sequence, ENST00000541893.5.

A

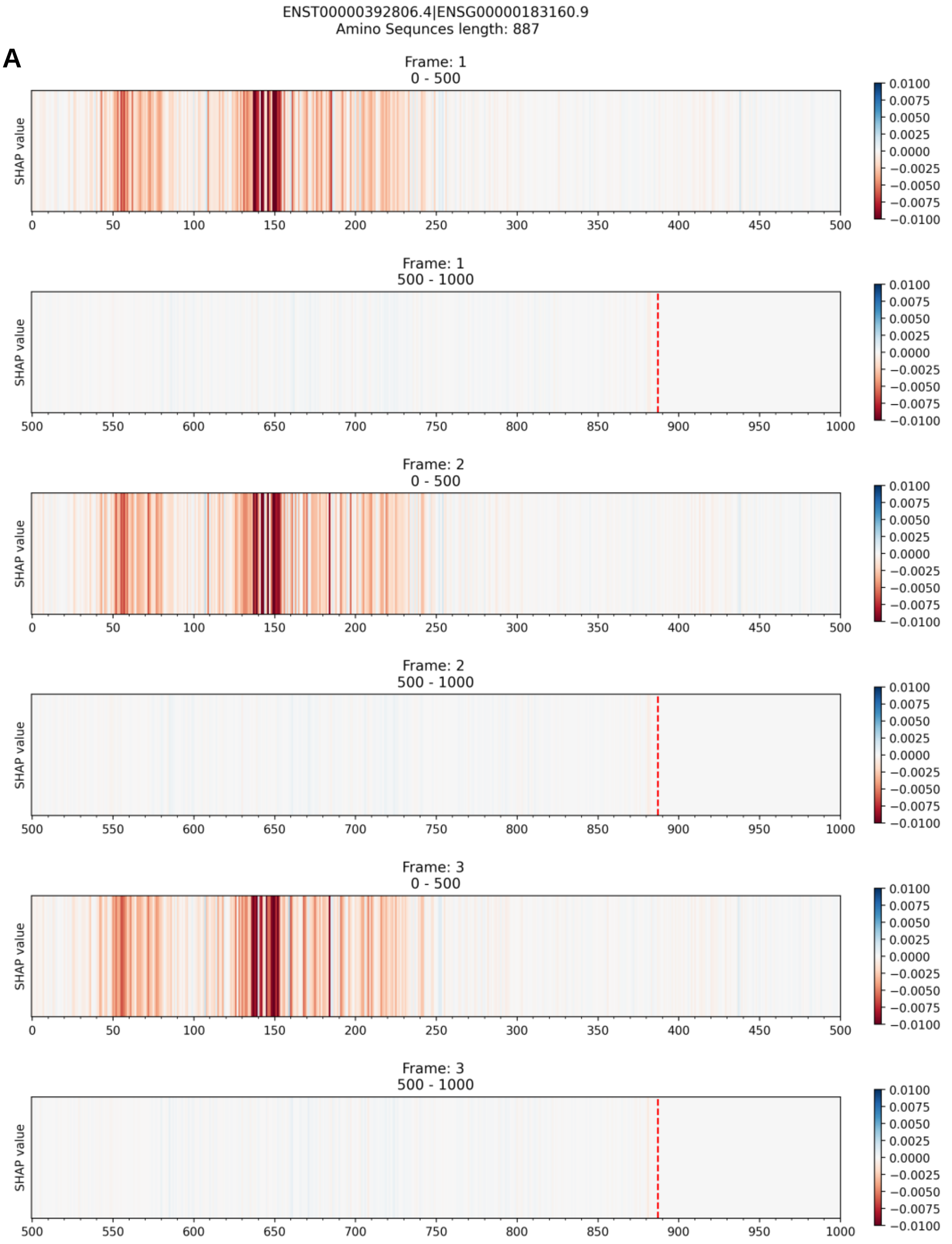


B

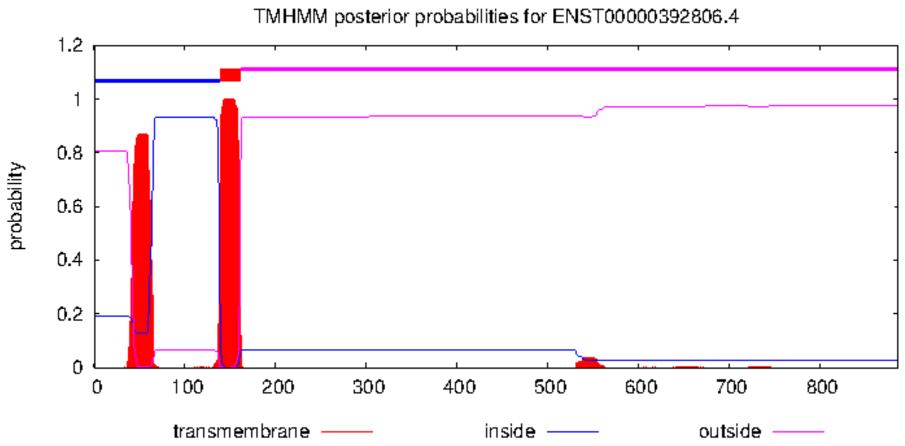


**Figure S10.** (**A**) The explanation result of Xlnc1DCNN and (**B**) the prediction result of the transmembrane helices by TMHMM program, on the true negative sequence, ENST00000392806.4.

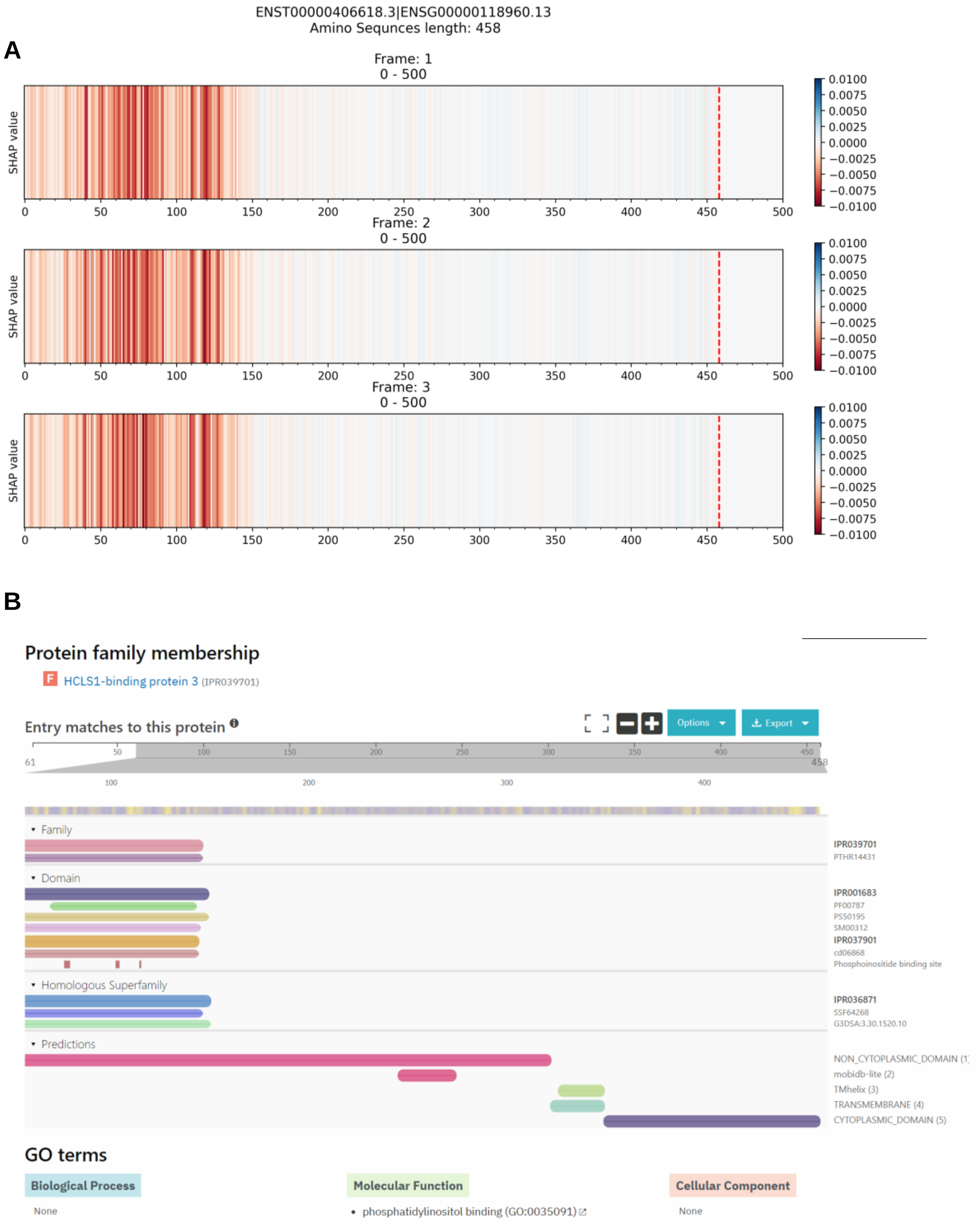


**Figure S11.** (**A**) The explanation result of Xlnc1DCNN and (**B**) the HCLS1-binding protein 3 (IPR039701) family and Phox homology (IPR001683) domain identified by InterPro, on the true negative sequence, ENST00000406618.3.

A

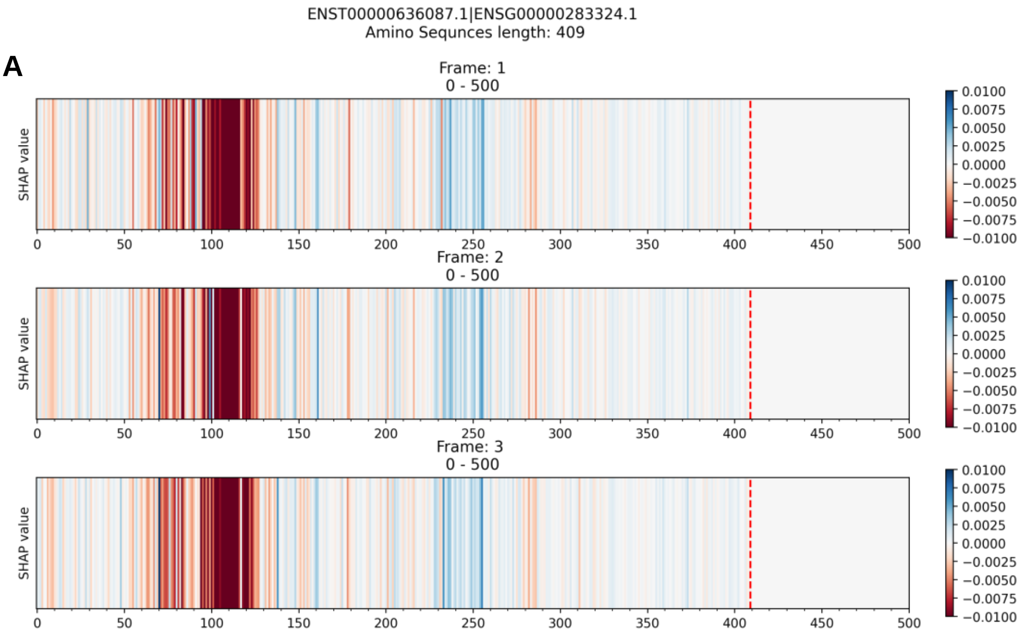


B

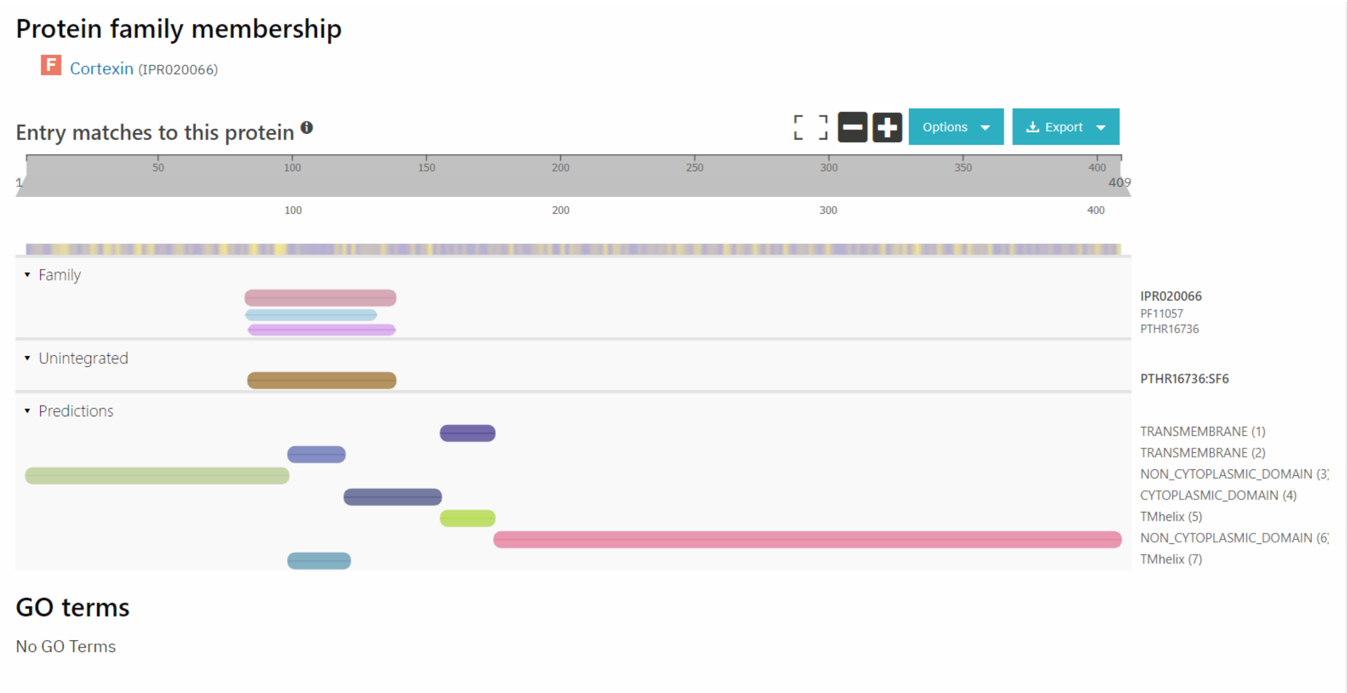


**Figure S12**. (**A**) The explanation result of Xlnc1DCNN and (**B**) the Cortexin (IPR020066) family identified by InterPro, on the true negative sequence, ENST00000636087.1.

A

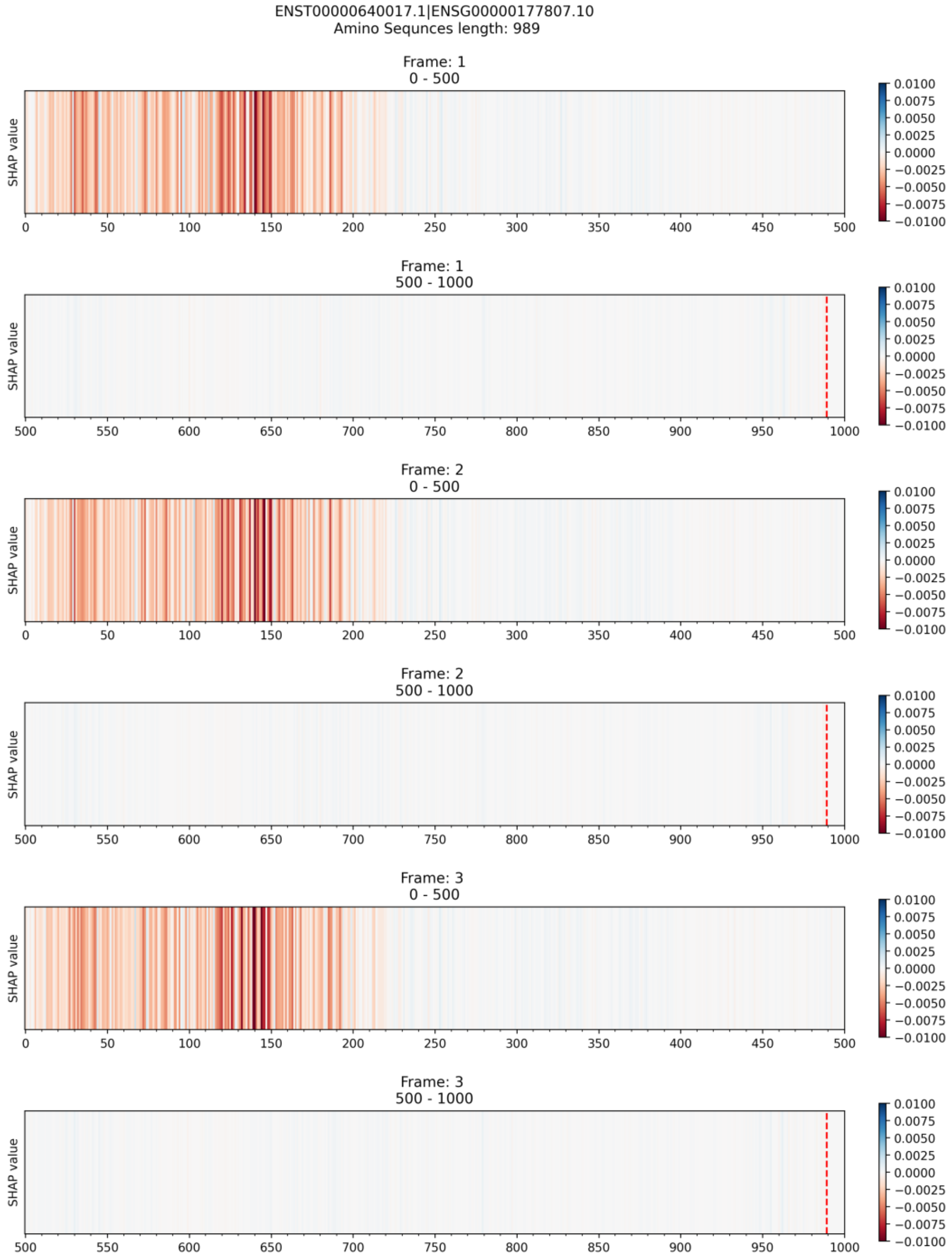
ENST00000640017.1|ENSG00000177807.10
Amino Sequnces length: 989
Frame: 1
0 - 500
Frame: 1
500 - 1000
Frame: 2
0 - 500
Frame: 2
500 - 1000
Frame: 3
0 - 500
Frame: 3
500 - 1000
SHAP value
0.0100
0.0075
0.0050
0.0025
0.0000
−0.0025
−0.0050
−0.0075
−0.0100

B

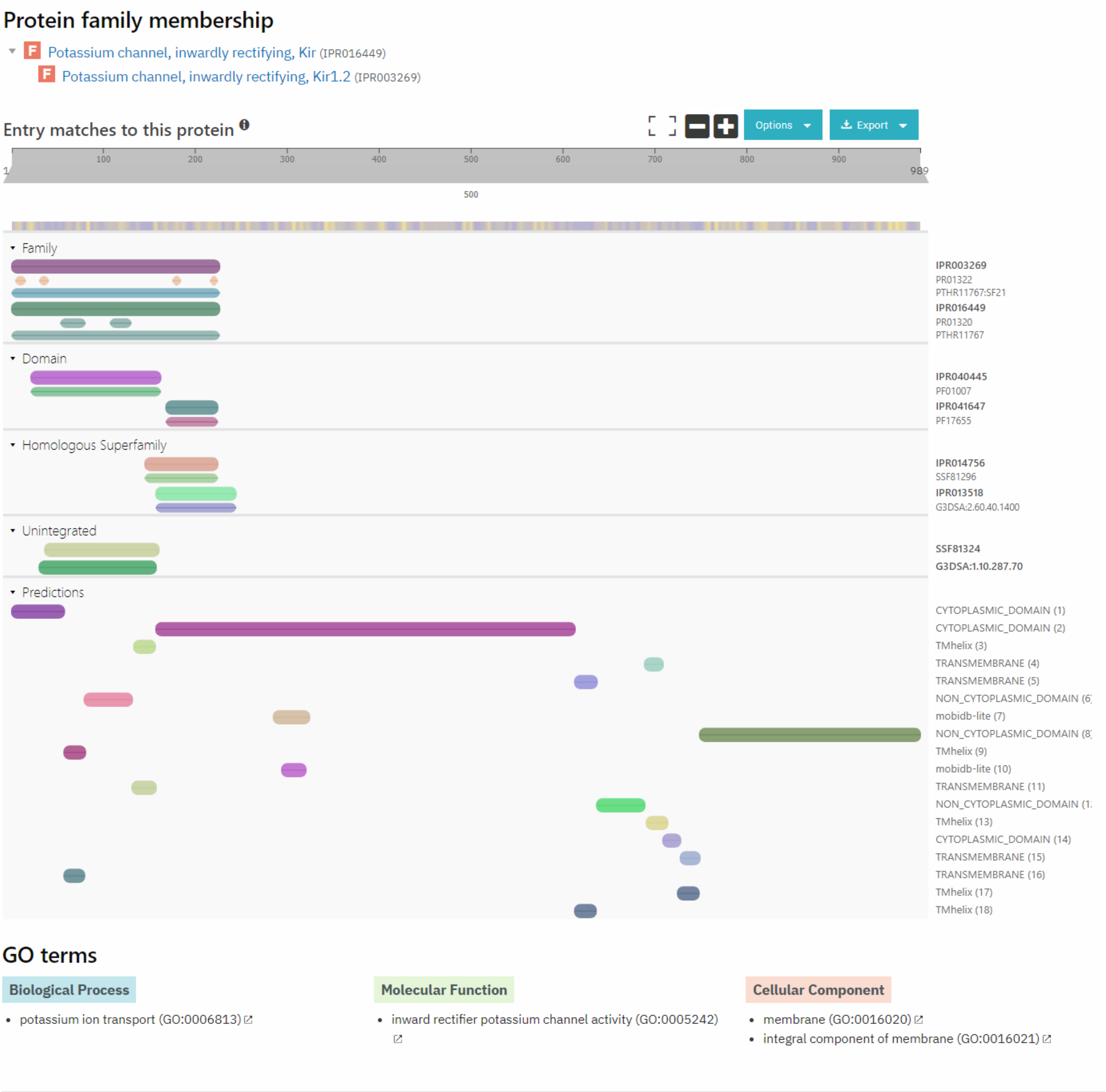


**Figure S13.** (A) The explanation result of Xlnc1DCNN and (B) the Potassium channel, inwardly rectifying, Kir (IPR016449), Potassium channel, inwardly rectifying, Kir1.2 (IPR003269) families and Potassium channel, inwardly rectifying, transmembrane domain (IPR040445), Inward rectifier potassium channel, C-terminal (IPR041647) domains identified by InterPro, on the true negative sequence, ENST00000640017.1.

A

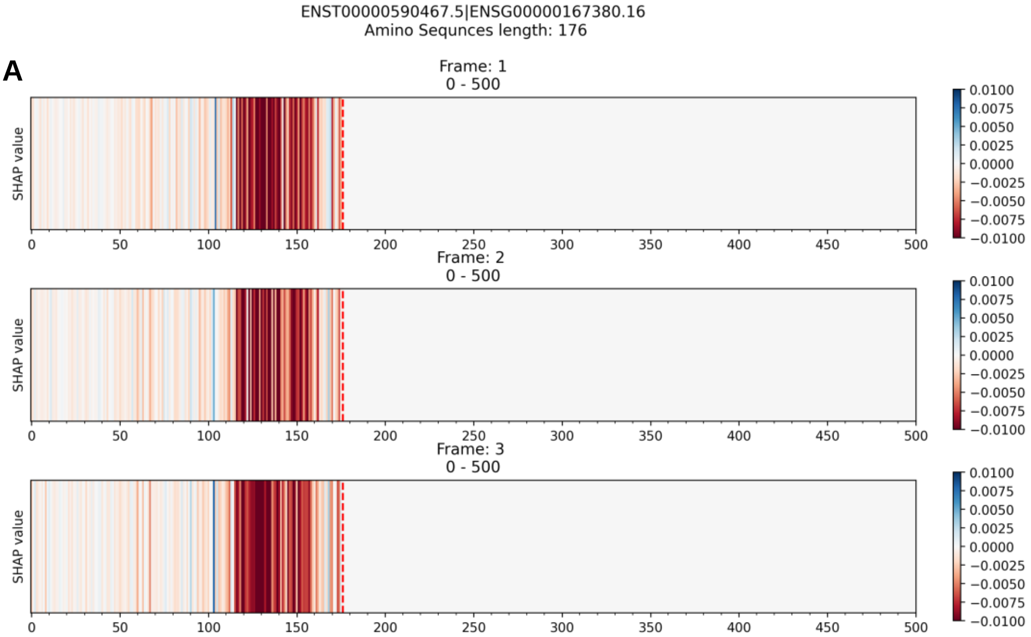


B

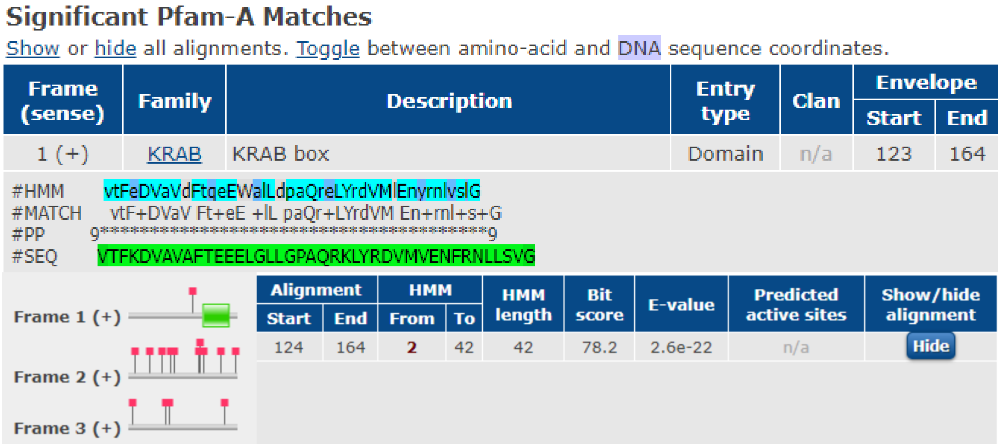

### Significant Pfam-A Matches

Show or hide all alignments. Toggle between amino-acid and DNA sequence coordinates.

| Frame (sense) | Family | Description | Entry type | Clan | Envelope | |
|---|---|---|---|---|---|---|
| | | | | | Start | End |
| 1 (+) | KRAB | KRAB box | Domain | n/a | 123 | 164 |

```
#HMM      vtFeDVaVdFtqeEWalLdpaQreLYrdVMlEnyrnlvslG
#MATCH    vtF+DVaV Ft+eE +lL paQr+LYrdVM En+rnl+s+G
#PP       9***************************************9
#SEQ      VTFKDVAVAFTEEELGLLGPAQRKLYRDVMVENFRNLLSVG
```

Frame 1 (+)

Frame 2 (+)

Frame 3 (+)

| Alignment | | HMM | | HMM length | Bit score | E-value | Predicted active sites | Show/hide alignment |
|---|---|---|---|---|---|---|---|---|
| Start | End | From | To | | | | | |
| 124 | 164 | 2 | 42 | 42 | 78.2 | 2.6e-22 | n/a | Hide |

**Figure S14.** (A) The explanation result of Xlnc1DCNN and (B) the KRAB domain identified by Pfam, on the true negative sequence, ENST00000590467.5.

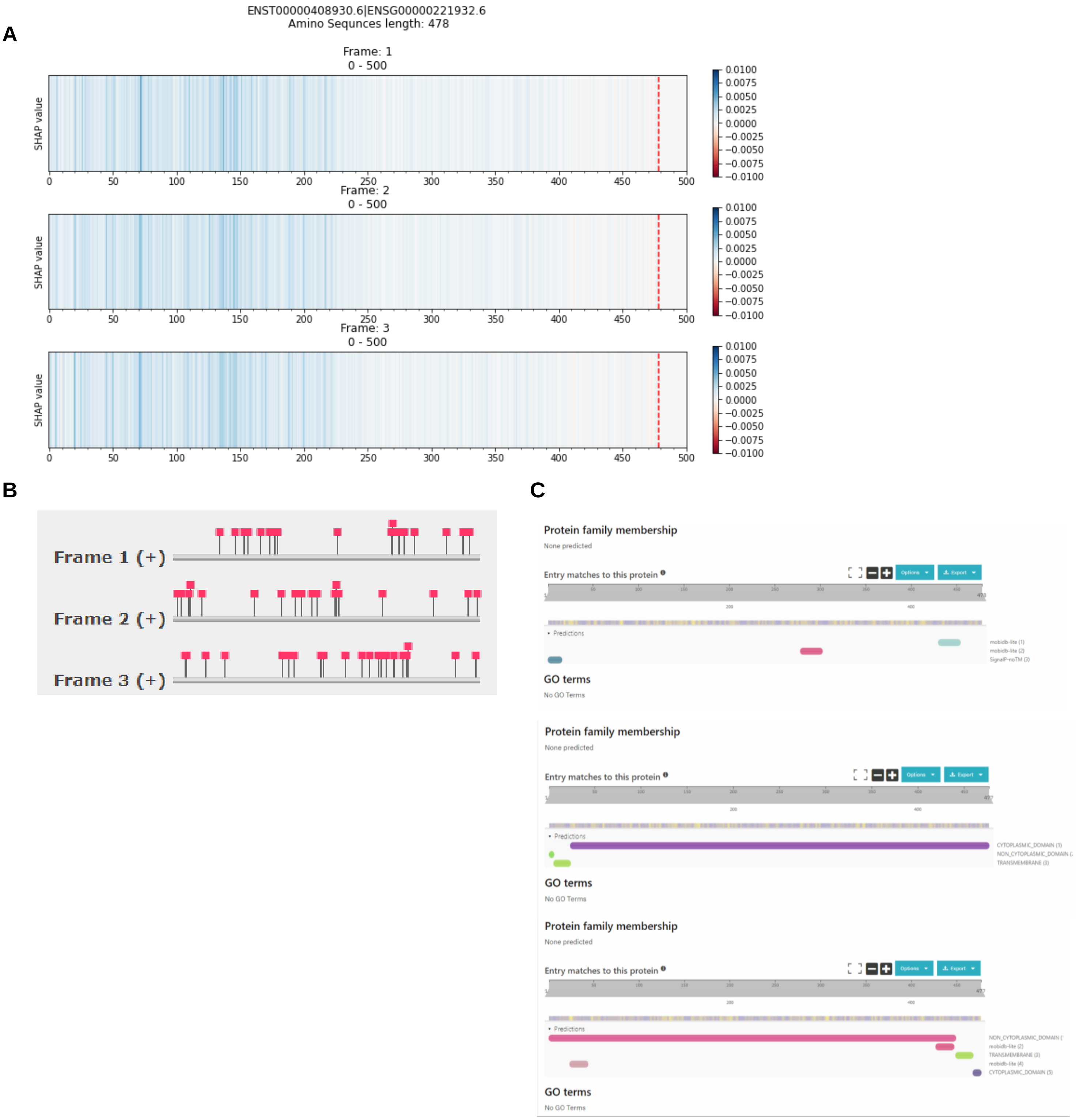


**Figure S15.** (**A**) The explanation result of Xlnc1DCNN, (**B**) The identification result from Pfam, and (**C**) The identification results from InterPro, which cannot identify any protein domains or families on the false positive sequence, ENST00000408930.6.

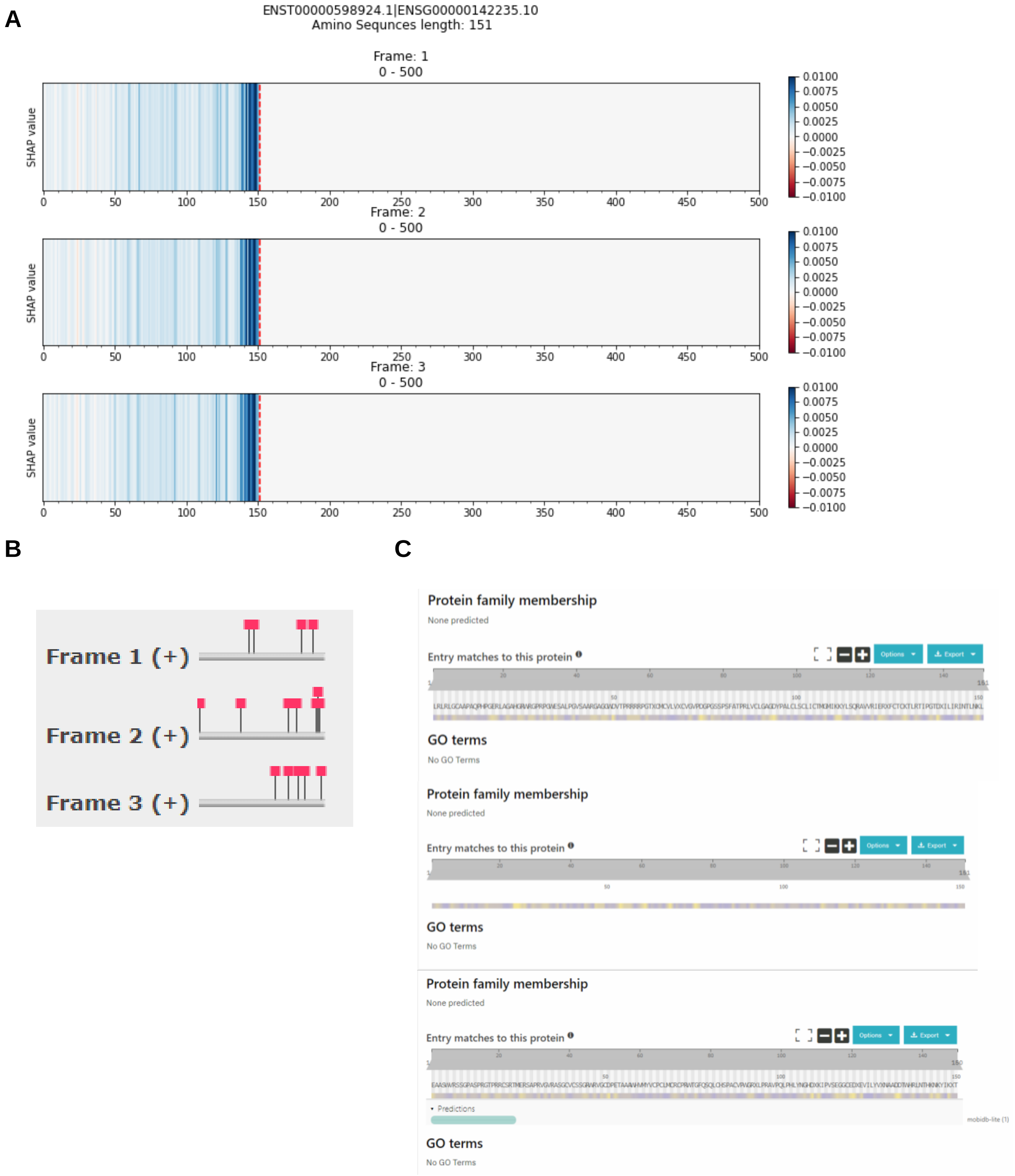


**Figure S16**. (**A**) The explanation result of Xlnc1DCNN, (**B**) The identification result from Pfam, and (**C**) The identification results from InterPro, which cannot identify any protein domains or families on the false positive sequence, ENST00000598924.1.

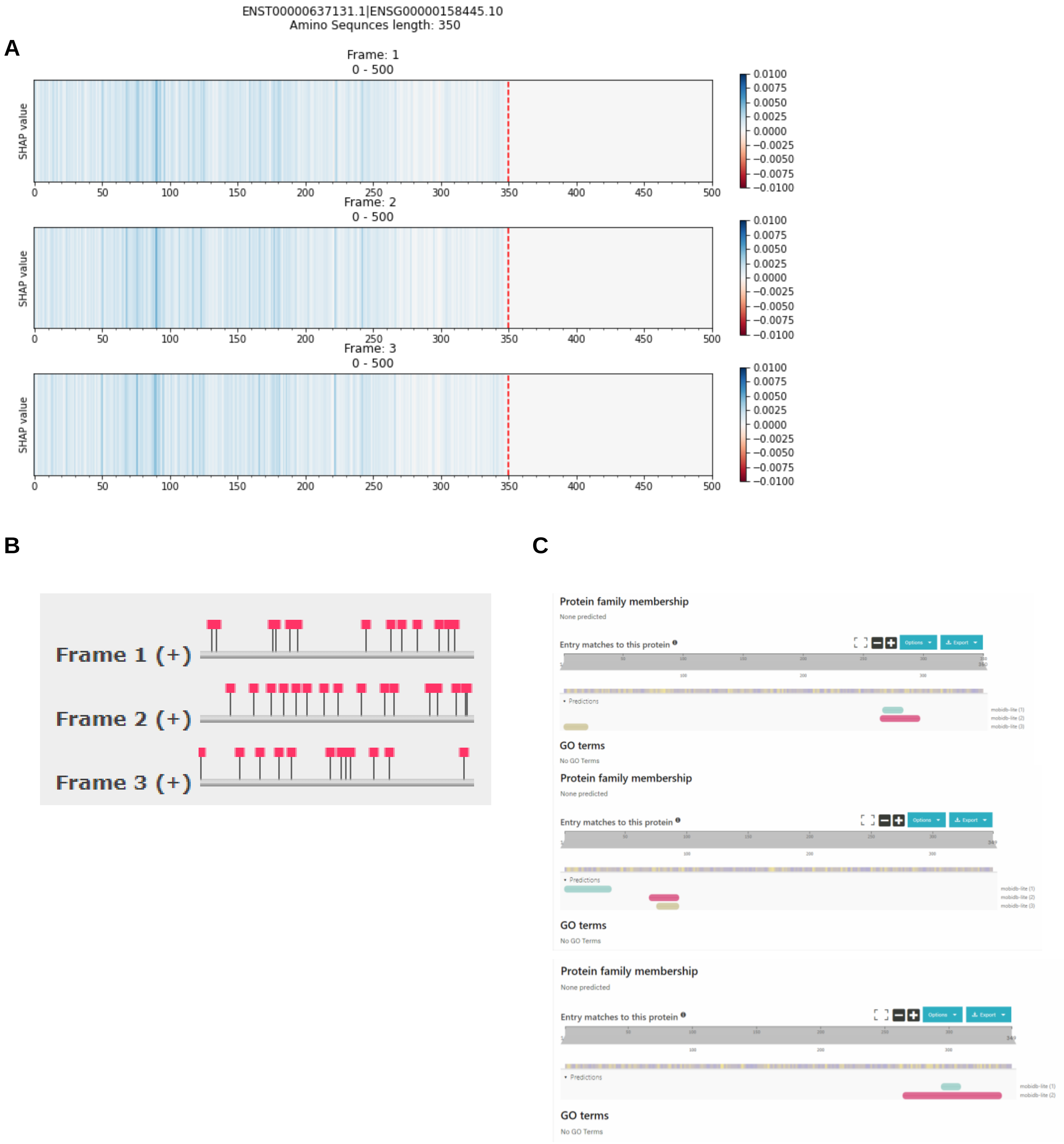


**Figure S17.** (**A**) The explanation result of Xlnc1DCNN, (**B**) The identification result from Pfam, and (**C**) The identification results from InterPro, which cannot identify any protein domains or families on the false positive sequence, ENST00000637131.1.

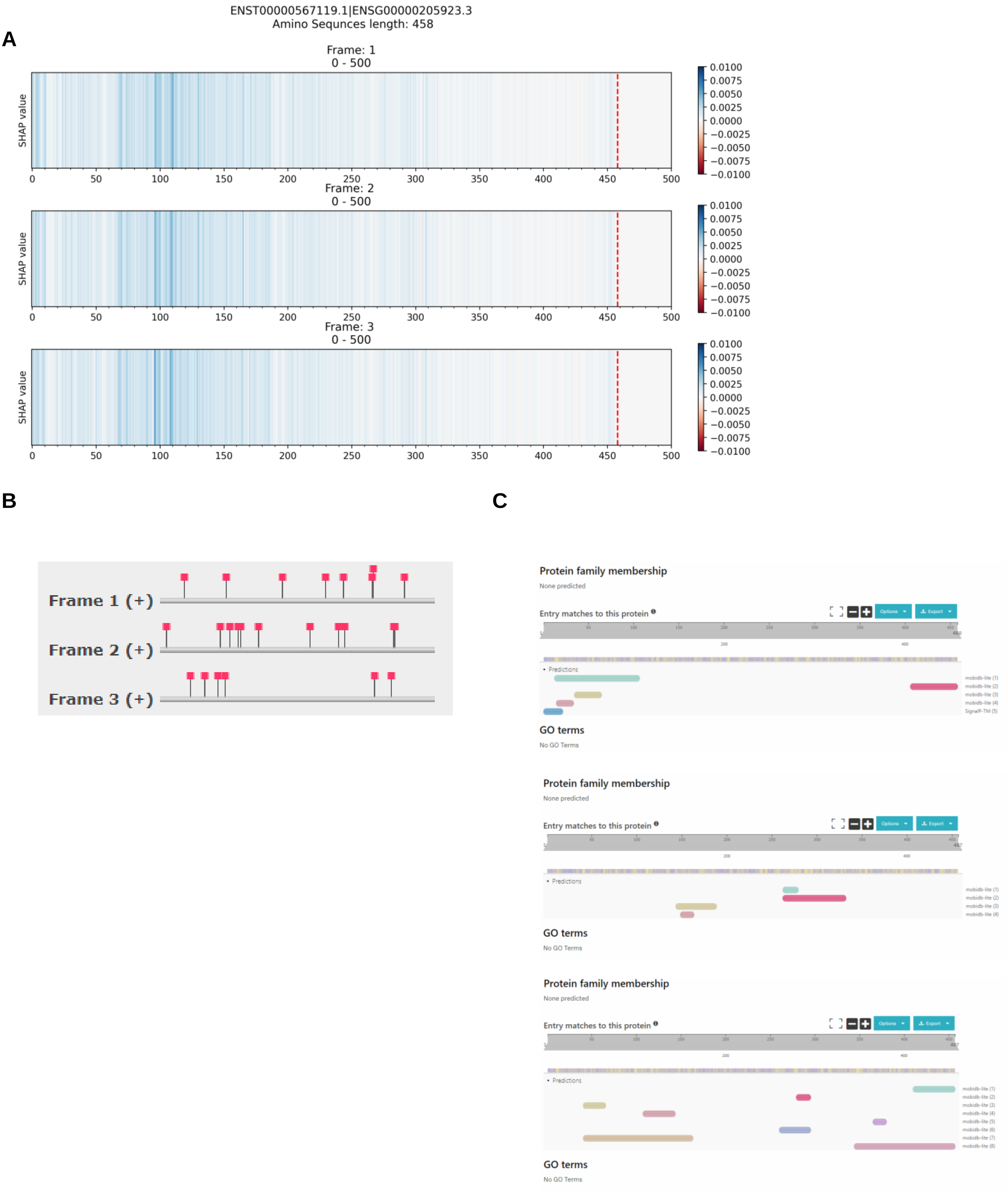


**Figure S18.** (**A**) The explanation result of Xlnc1DCNN, (**B**) The identification result from Pfam, and (**C**) The identification results from InterPro, which cannot identify any protein domains or families on the false positive sequence, ENST00000567119.1.

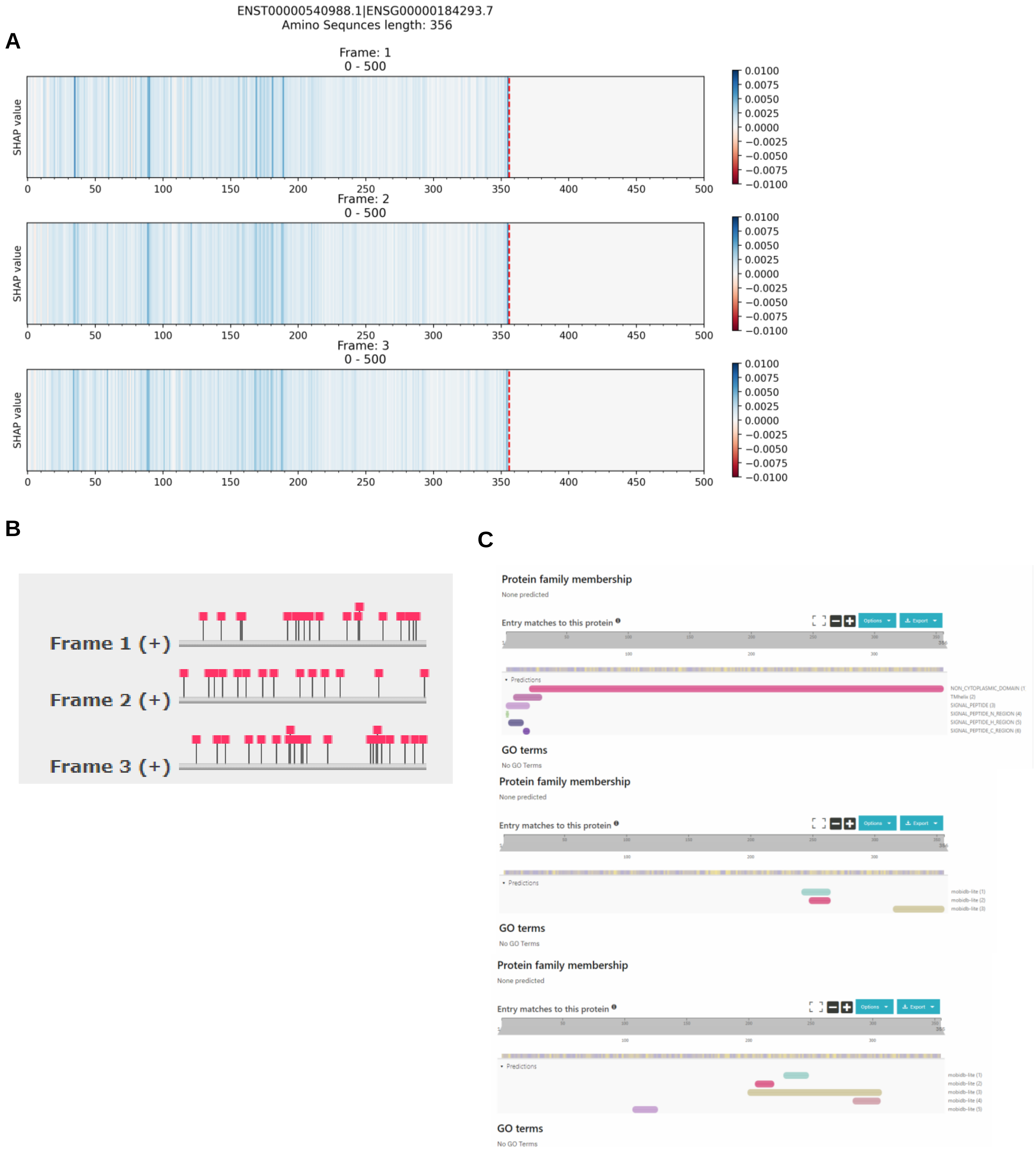


**Figure S19.** (**A**) The explanation result of Xlnc1DCNN, (**B**) The identification result from Pfam, and (**C**) The identification results from InterPro, which cannot identify any protein domains or families on the false positive sequence, ENST00000540988.1.

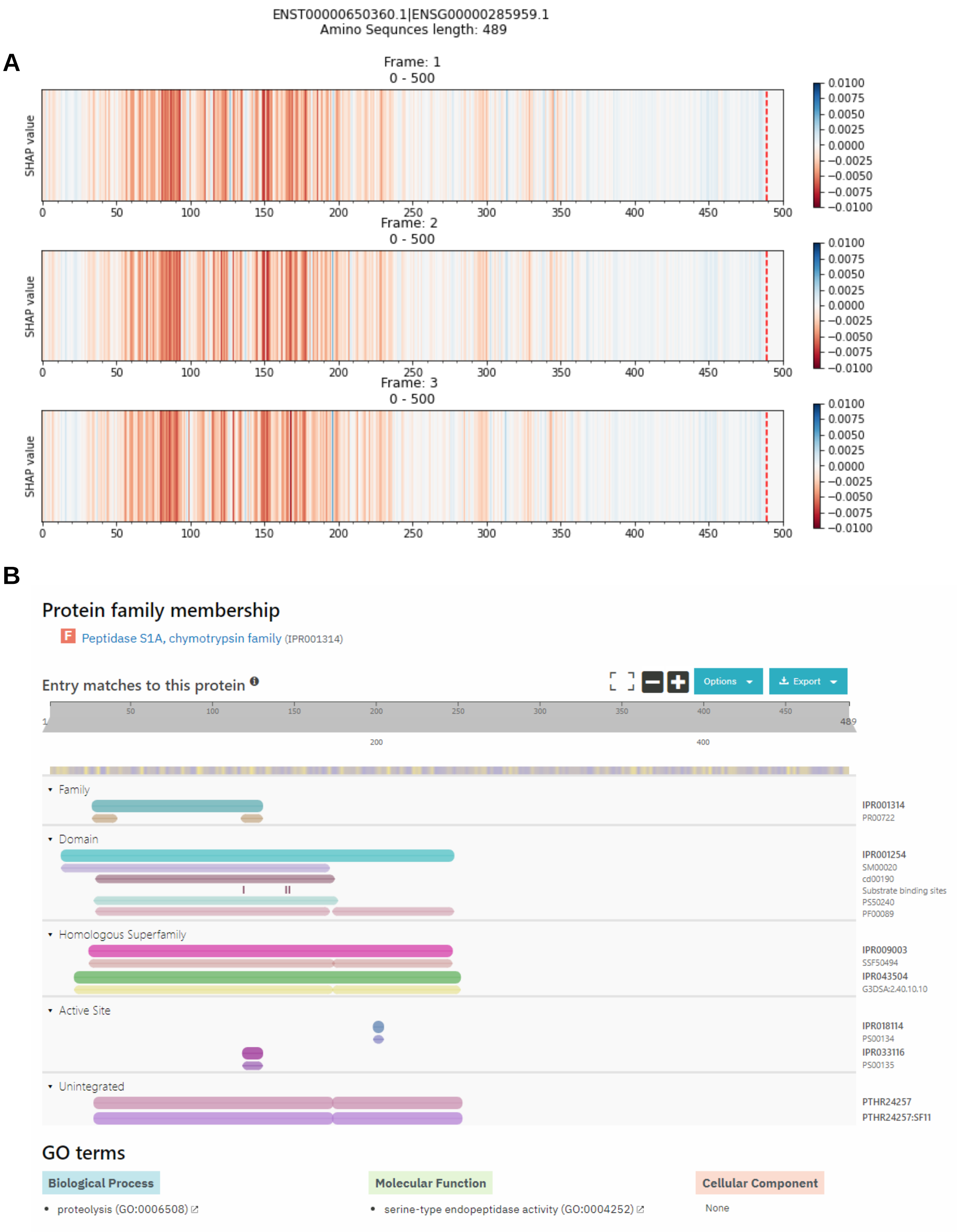


**Figure S20.** (**A**) The explanation result of Xlnc1DCNN and (**B**) the Peptidase S1A, chymotrypsin family (IPR001314) and Serine proteases, trypsin domain (IPR001254) domain identified by InterPro, on false negative sequence, ENST00000650360.1.

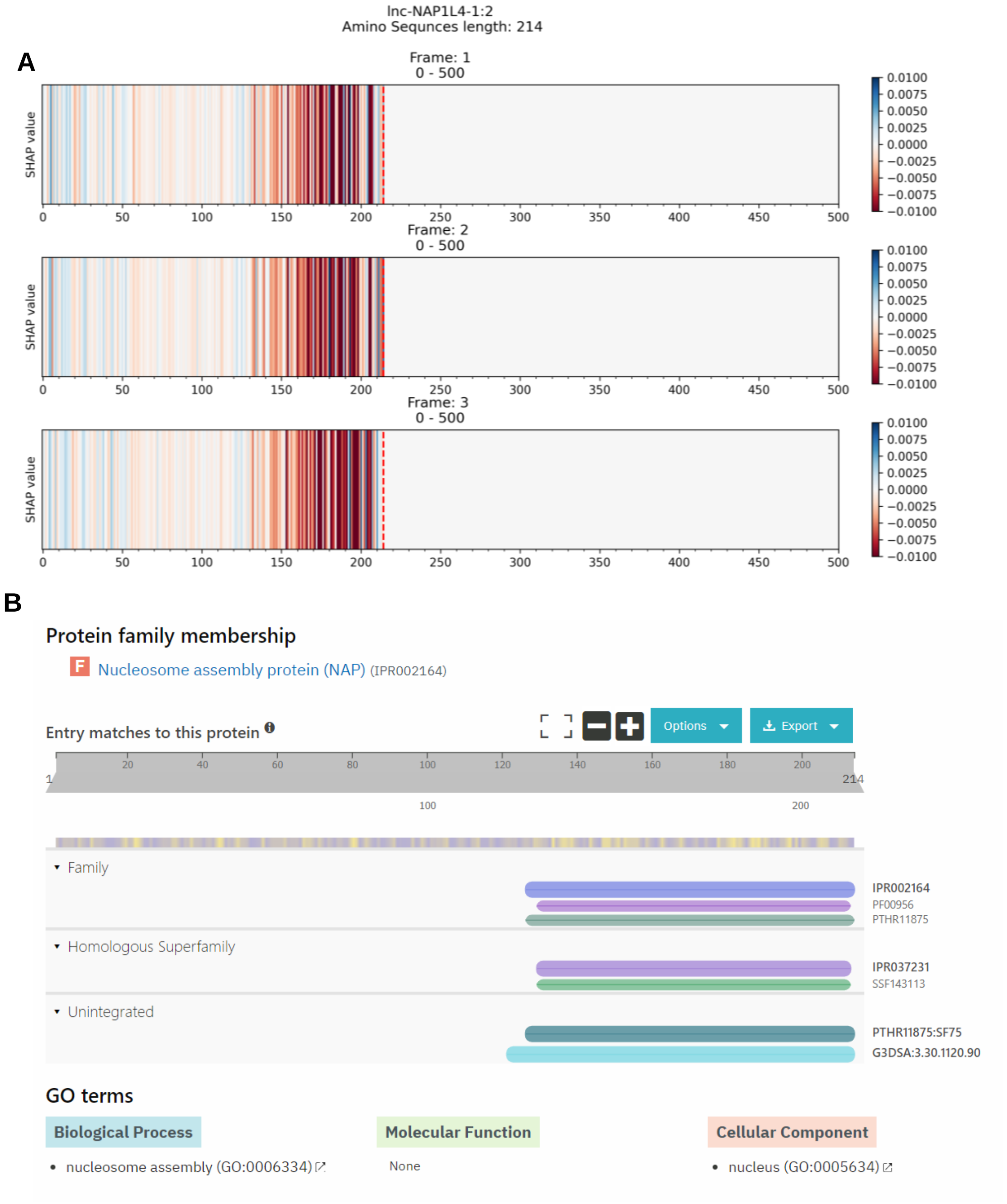


**Figure S21.** (**A**) The explanation result of Xlnc1DCNN and (**B**) the Nucleosome assembly protein (NAP) (IPR002164) family identified by InterPro, on false negative sequence, lnc-NAP1L4-1:2.

A

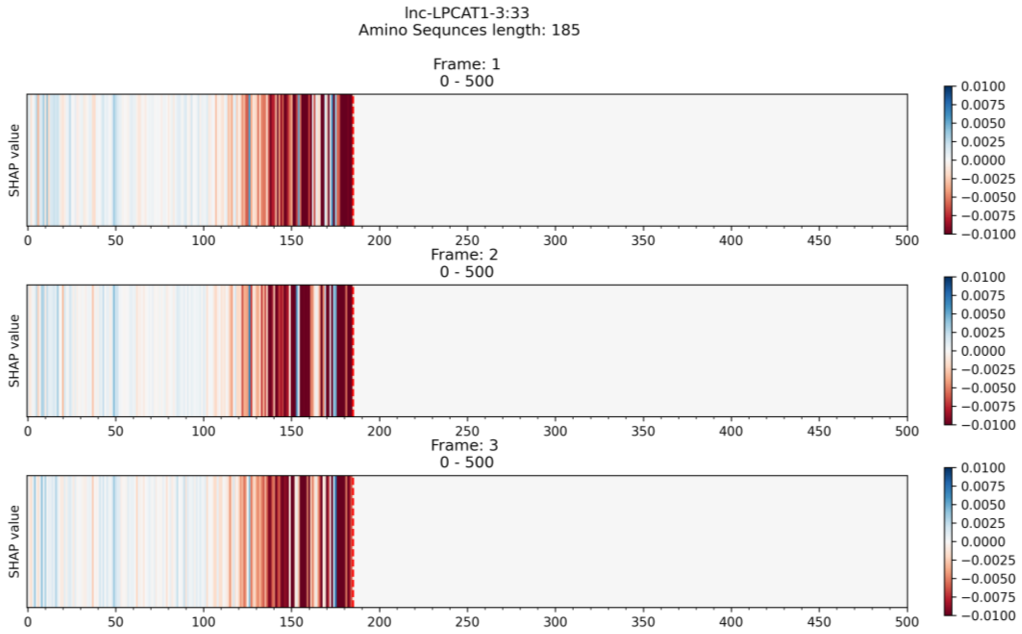


B

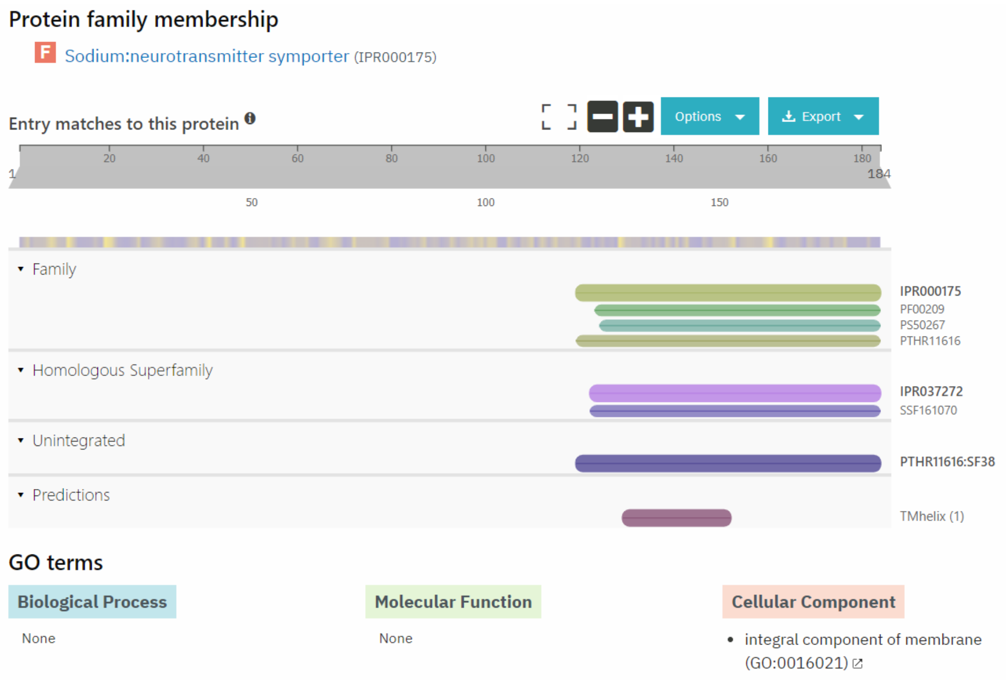


**Figure S22.** (**A**) The explanation result of Xlnc1DCNN and (**B**) the Sodium:neurotransmitter symporter (IPR000175) family identified by InterPro, on false negative sequence, lnc-LPCAT1-3:33.

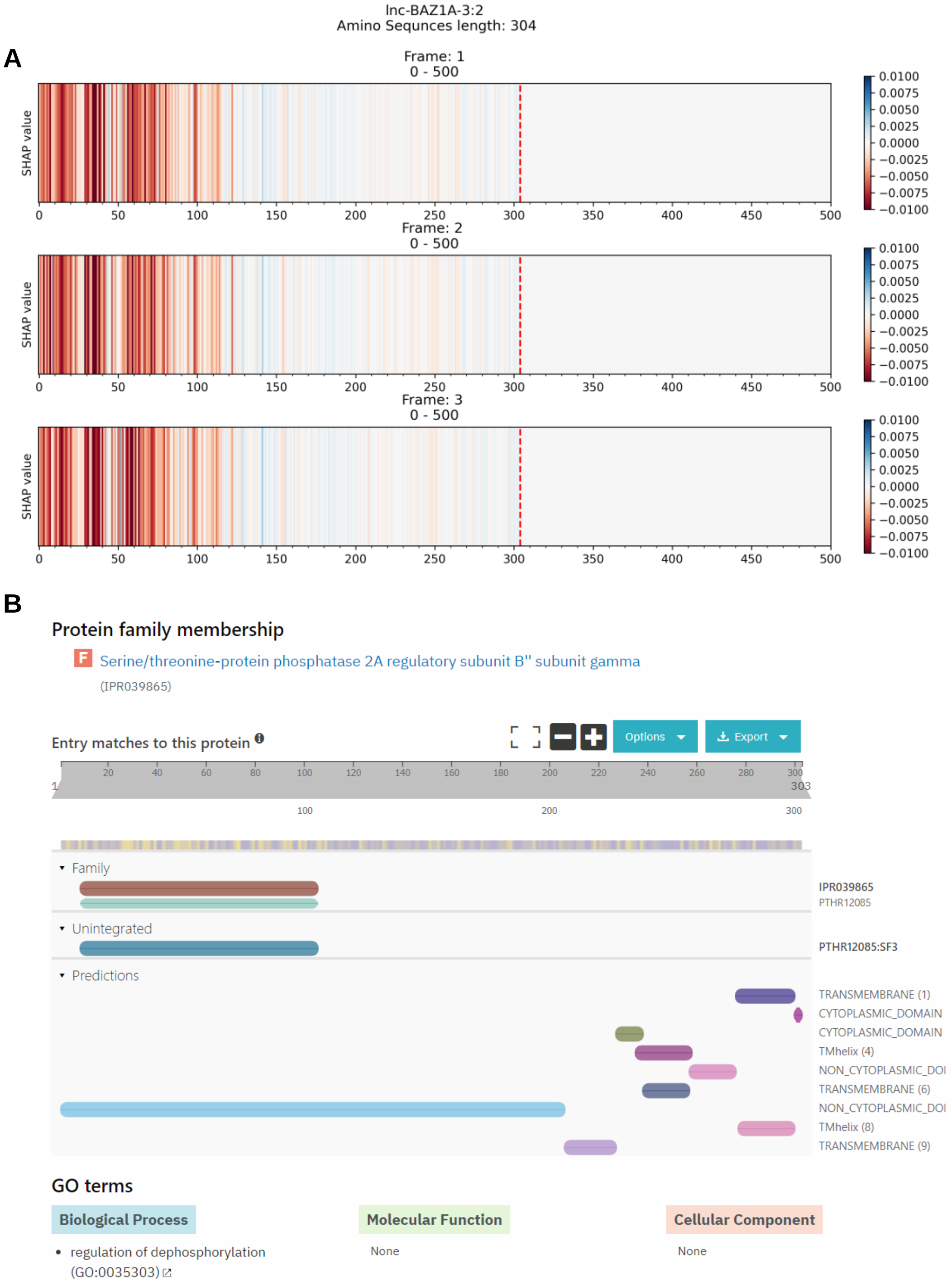


**Figure S23.** (**A**) The explanation result of Xlnc1DCNN and (**B**) the Serine/threonine-protein phosphatase 2A regulatory subunit B'' subunit gamma (IPR039865) family identified by InterPro, on false negative sequence, lnc-BAZ1A-3:2.

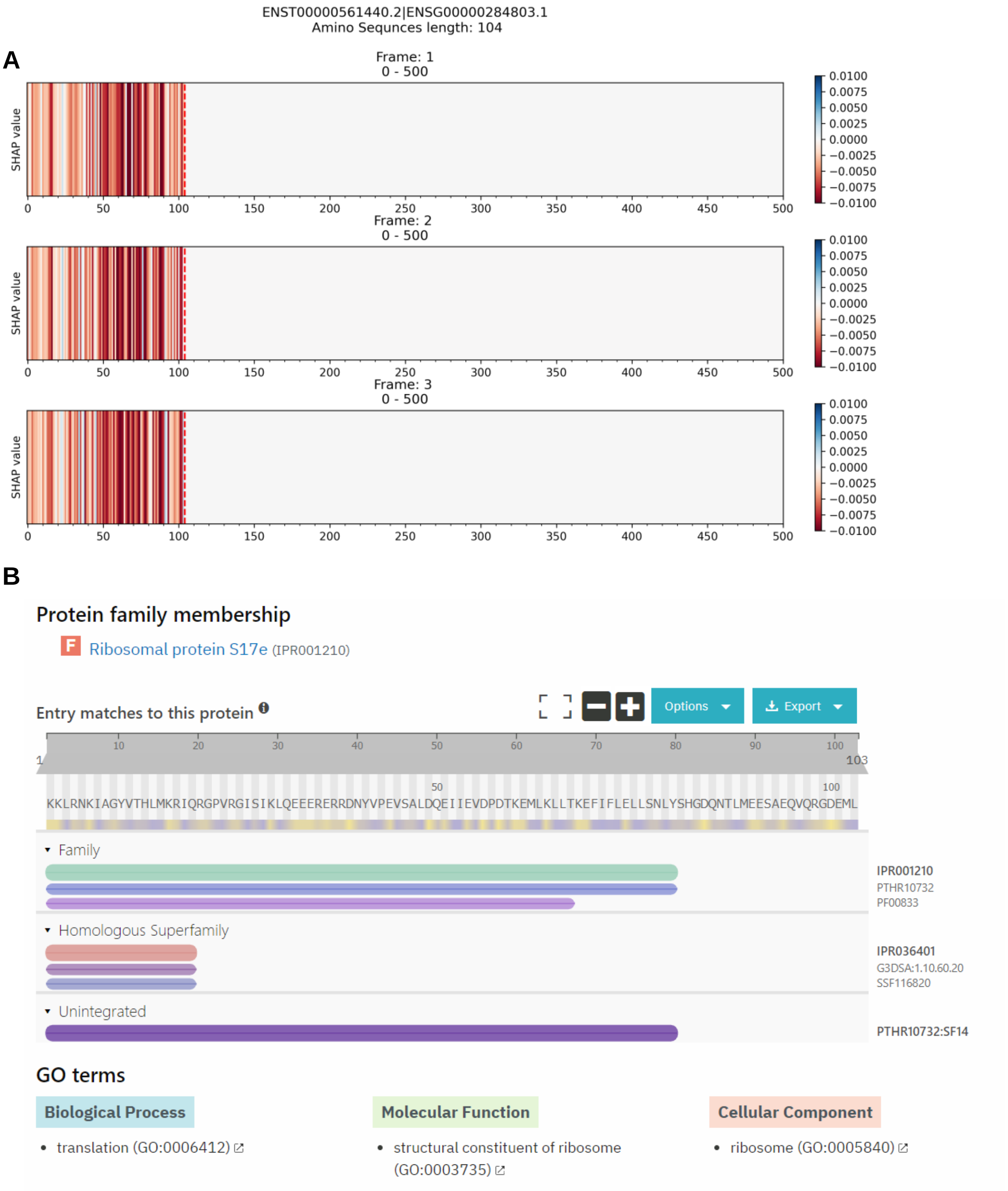


**Figure S24.** (**A**) The explanation result of Xlnc1DCNN and (**B**) the Ribosomal protein S17e (IPR001210) family identified by InterPro, on false negative sequence, ENST00000561440.2.

A

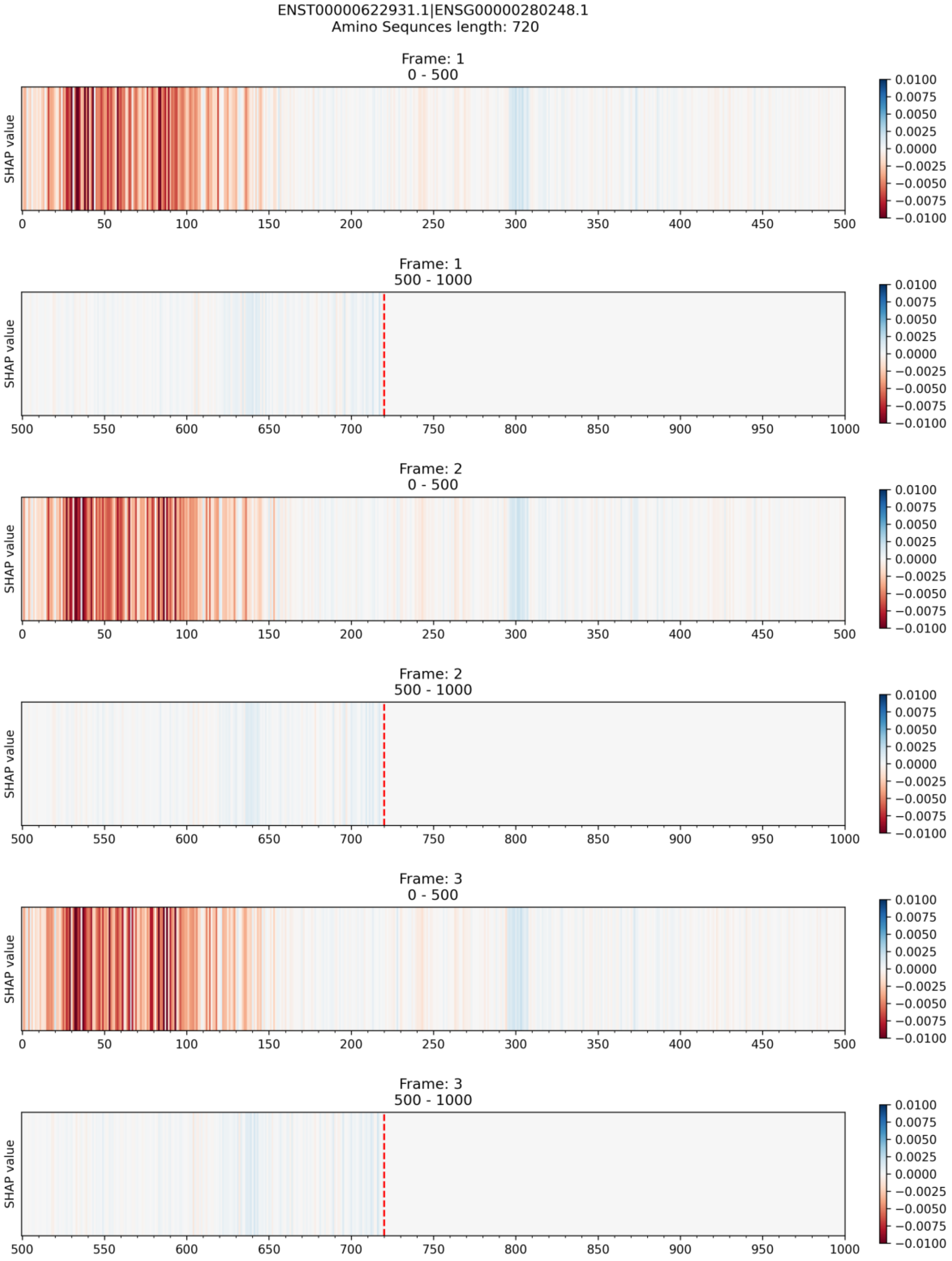
ENST00000622931.1|ENSG00000280248.1
Amino Sequnces length: 720
Frame: 1
0 - 500
Frame: 1
500 - 1000
Frame: 2
0 - 500
Frame: 2
500 - 1000
Frame: 3
0 - 500
Frame: 3
500 - 1000
SHAP value
0.0100
0.0075
0.0050
0.0025
0.0000
−0.0025
−0.0050
−0.0075
−0.0100

**B**

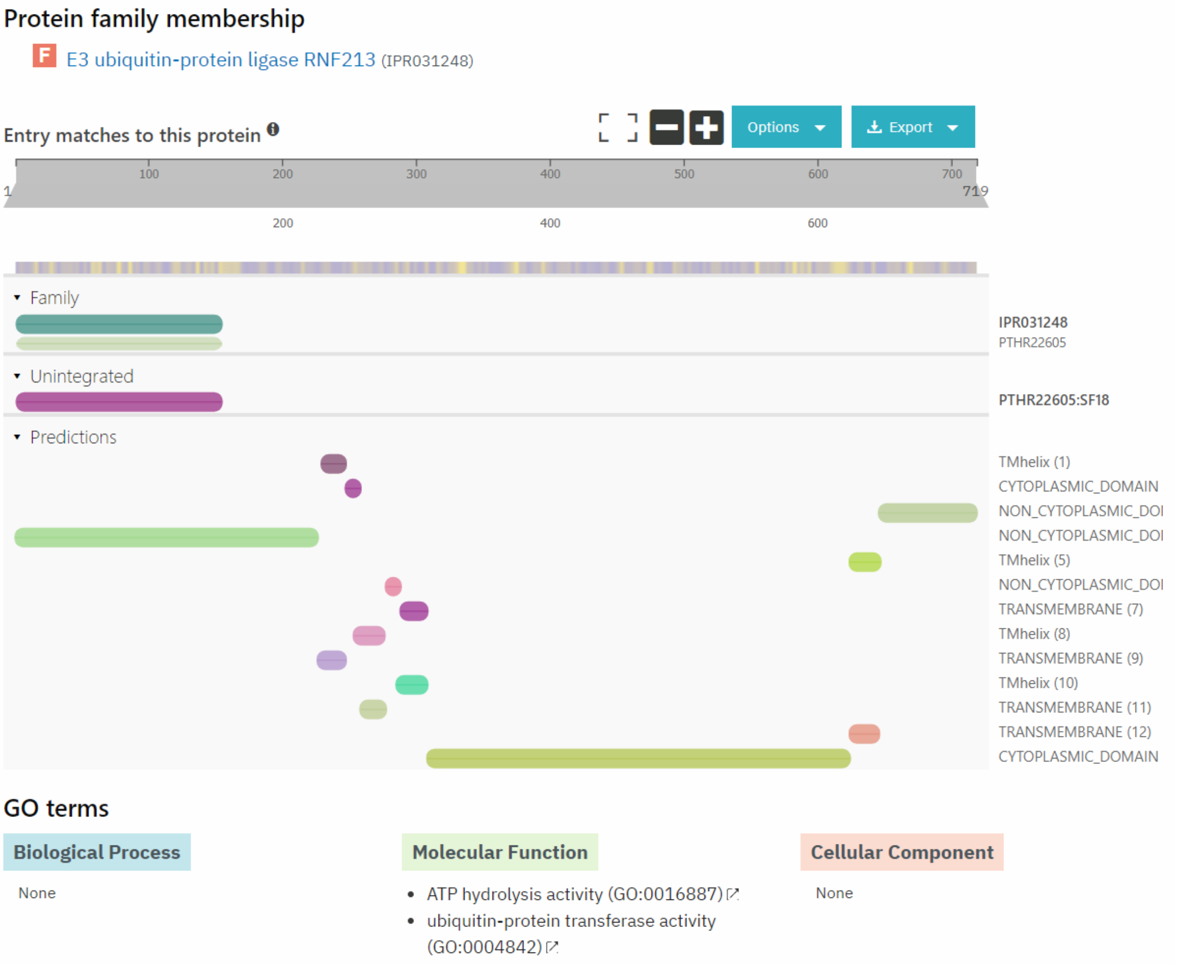


**Figure S25.** (**A**) The explanation result of Xlnc1DCNN and (**B**) the E3 ubiquitin-protein ligase RNF213 (IPR031248) family identified by InterPro, on false negative sequence, ENST00000622931.1.
